# Two Stages of Dynamic Metabolic and Transcriptomic Remodeling During the Adaptation to Caloric Restriction in Male C57BL/6J

**DOI:** 10.64898/2026.03.03.709429

**Authors:** Heidi H. Pak, Emil S. Rassmussen, Lauren Palluth, Joseph S. Takahashi

## Abstract

The molecular basis of caloric restriction (CR) has been defined primarily at a metabolic steady state, leaving the initiating events that drive the transition from *ad libitum* feeding to an adapted CR state largely unresolved. Here, we combine continuous indirect calorimetry with longitudinal bulk RNA-seq of liver and inguinal white adipose tissue (iWAT) sampled at six circadian timepoints across four stages of adaptation to 30% CR in male C57BL/6J mice. We show that whole-body metabolic adaptation proceeds through two discrete adaptive phases separated by a threshold at approximately 14 days; during this initial transition, consolidated feeding attenuates ketogenesis, establishing a distinct whole-body metabolic phenotype prior to long-term adaptation. To elucidate the molecular mechanisms underlying these physiological shifts, weighted gene co-expression network analysis (WGCNA) was performed, revealing that hepatic transcriptional remodeling is organized proportionally to fasting duration, whereas iWAT remodeling remains restricted to specific circadian timepoints. Because systemic adaptation requires coordinated inter-tissue communication, we conducted a cartographic analysis to evaluate network topology and inter-modular connectivity. This approach identifies restricted populations of early kinless and connector hub genes, nucleated by *Casp3* in the liver and *Lpl* in iWAT, whose structural integration is established prior to the broader transcriptional remodeling observed at later timepoints. Functional annotation indicates the hepatic hub network is enriched for mitochondrial bioenergetics and FOXO/TP53-mediated transcription, while the iWAT hub network exhibits a bifurcated enrichment spanning ribosomal biosynthesis and immune-regulatory signaling. Although these tissues exhibit distinct transcriptional profiles, projecting both datasets onto a shared phenotypic eigenspace reveals a unified systemic response; as CR is maintained, dynamically regulated transcripts in both liver and iWAT converge on an adiponectin-coupled state. Ultimately, the identification of adiponectin as an integrative signal coordinating chronic adaptation across metabolically distinct tissues delineates the temporal sequence of early CR adaptation; furthermore, it establishes a mechanistic framework defining how early molecular and physiological shifts converge to achieve steady-state metabolic homeostasis.

## Introduction

Nearly a century has passed since the first experimental observations that reducing food intake could dramatically extend the lifespan of laboratory rodents, and calorie restriction (CR) remains, to this day, the most robust and reproducible intervention for promoting longevity and metabolic health across a diverse range of model organisms^1–6^. In the decades that followed those early discoveries, the field has made remarkable progress in identifying the metabolic effects of CR: reduced adiposity, improved insulin sensitivity, decreased systemic inflammation, enhanced mitochondrial efficiency, and protection against a broad spectrum of age-related diseases^7–12^. Yet despite this wealth of biological insight, a fundamental question remains unanswered: how does eating less translate into living longer and healthier?

The search for a mechanistic answer has implicated nearly every major nutrient-sensing and energy-regulating pathway; mTORC1 inhibition, AMPK activation, sirtuin-dependent deacetylation, insulin/IGF-1 signaling attenuation, and circadian rhythms have each been proposed as primary mediators of the CR response^13–20^. Individual manipulation of any one of these pathways can recapitulate select features of CR, lending credibility to each as a candidate mechanism, yet none has emerged as a unifying explanation. This is perhaps unsurprising; CR fundamentally reorganizes the relationship between an organism and its energy supply, and in doing so, simultaneously engages dozens of interconnected regulatory networks across multiple tissues. The result is a system so comprehensively remodeled that distinguishing causal drivers from downstream consequences becomes extraordinarily difficult.

This challenge is compounded by the redundant nature of the signaling networks involved. When one pathway is ablated, others compensate, buffering the system against perturbation and obscuring the contribution of any individual node^15,21–28^. Traditional loss-of-function approaches, powerful in simpler biological contexts, struggle to resolve causal hierarchy in a network this deeply interconnected. The field has thus reached a paradoxical position: the more we learn about the molecular effects of CR, the harder it becomes to identify which of these effects actually matter.

One approach to cut through this complexity is to shift the question. Rather than asking what differs between a CR animal and an ad libitum-fed control at steady state, a comparison that captures the cumulative endpoint of adaptation rather than its mechanistic origin, we can ask instead – what changes first? If CR acts through a primary regulatory event that subsequently drives the reorganization of downstream networks, that event should be detectable as an early, persistent, and highly connected node in the transcriptional landscape, one that precedes and predicts the broader adaptive response. Capturing this initiating signal requires studying animals not after they have fully adapted to CR, but during the transition itself.

Existing transcriptomic studies of CR have not been designed with this question in mind. Most analyses have been performed well after the establishment of a CR steady state^29,30^, and tissues have typically been sampled at a single time of day^31–34^, collapsing the rich circadian regulation of gene expression into a static snapshot. Similarly, while time-restricted feeding (TRF) paradigms have been extensively studied, these models do not recapitulate the consolidated feeding pattern and extended daily fast that define CR in rodents^35–37^, leaving the question of how CR specifically restructures the tissue-level metabolic architecture, across both the adaptive timeline and during the circadian cycle, remains largely unaddressed.

To address this gap, we designed a longitudinal study to capture the metabolic response and remodeling of the transcriptome as mice transitioned from *ad libitum* feeding to 30% CR over a four-week period. Continuous indirect calorimetry revealed that whole-body metabolic adaptation unfolds through two discrete stages, with the major transition threshold emerging at approximately two weeks. This finding provided a principled, physiology-driven framework for subsequent tissue collection: liver and inguinal white adipose tissue were sampled at six circadian timepoints across four stages of adaptation, spanning early adaptation, late adaptation, and the established metabolic steady state. By mapping the transcriptional landscape of adaptation as it unfolds rather than after it has settled, this study provides a framework for identifying the early, persistent, and highly connected regulatory nodes that may mediate the systemic transition to the CR state and offers a new lens through which the mechanistic basis of CR’s broad metabolic benefits can be examined.

## Results

### Continuous indirect calorimetry reveals two discrete phases of metabolic adaptation to caloric restriction

To capture the kinetics of systemic adaptation to sustained negative energy balance, we housed male C57BL/6J mice in continuous indirect calorimetry chambers over a four-week period (**Fig. 1**). This approach enabled simultaneous, high-resolution tracking of whole-body energy expenditure (EE), respiratory exchange ratio (RER), oxygen consumption (VO2), carbon dioxide production (VCO2), and locomotor activity. Mice were acclimated under ad libitum (AL) conditions for 48 hours to establish baseline metabolic profiles prior to CR initiation. Food was removed on the morning before the intervention began, and 30% CR was introduced at ZT12, imposing an 8-hour daytime fast before the first restricted meal.

**Figure 1.**
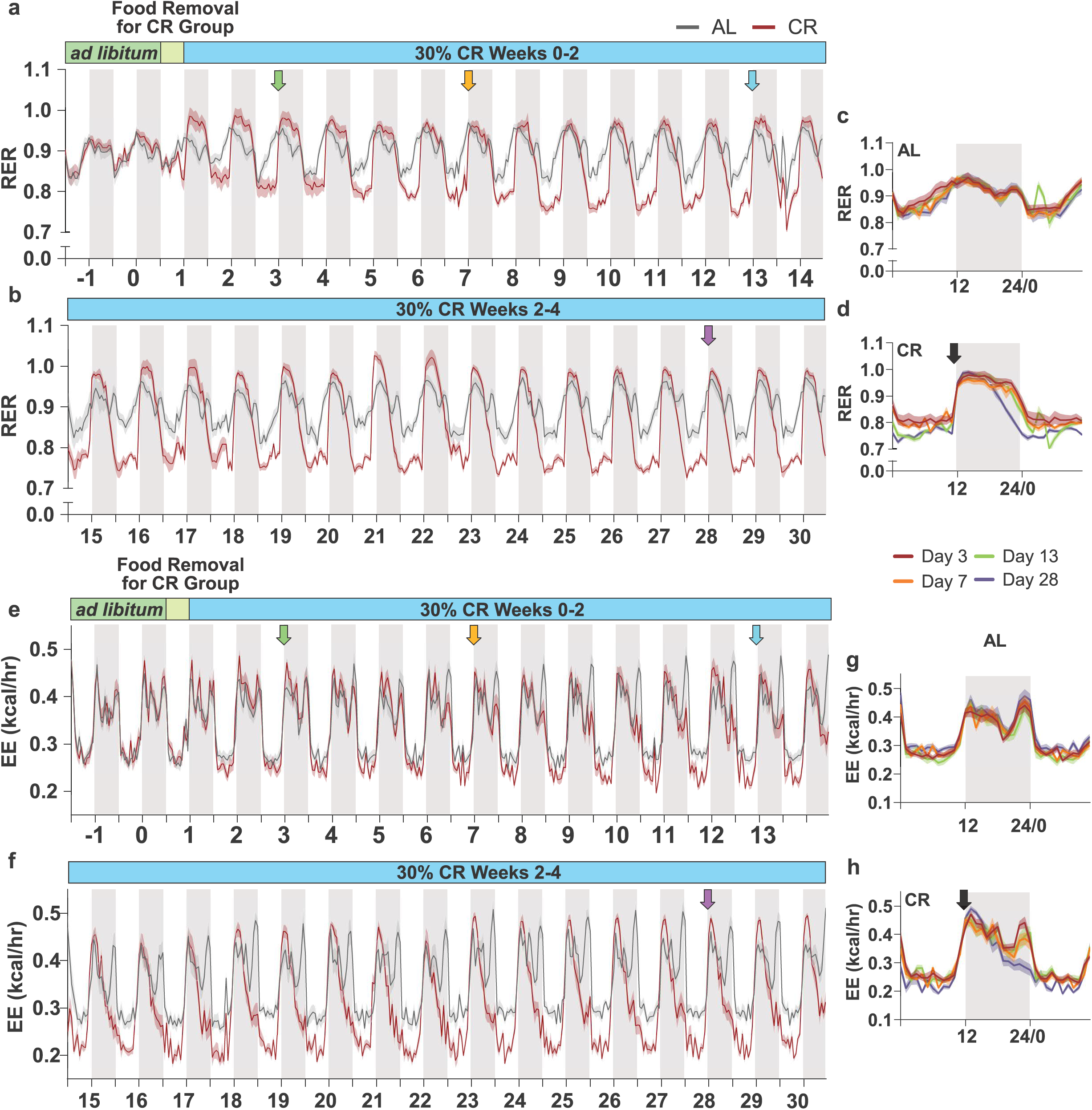
Continuous indirect calorimetry reveals a two-week threshold for systemic metabolic adaptation to caloric restriction. 4 week time course recording of respiratory exchange ratio (RER) and energy expenditure (EE). All animals were acclimated for 48 hrs on *ad libitum conditions,* fasted for ∼8 hours prior to the initiation of CR, and placed on CR for four weeks. **a, b,** RER versus time. **c, d,** Overlay of longitudinal RER profiles for AL and CR animals at Days 3, 7, 13, and 28. **e, f,** Energy expenditure (EE) versus time. **g, h** Overlay of longitudinal RER profiles for AL and CR animals at Days 3, 7, 13, and 28. Data are presented as mean ± SEM. n = 10-12 per group. Shaded regions denote the dark cycle. Colored arrows in **a, b, e, f** represent the timing for **c, d, g, h.** Black arrows represent the time when CR animals were fed. Data are mean ± s.e.m.; n = 10-12 per feeding condition.

During acclimation, all animals exhibited consistent diurnal RER patterns (**Fig. 1a**). Upon CR initiation, RER values rose promptly to ∼1.0, reflecting rapid carbohydrate oxidation within the first day of restriction (Fig. 1a). This early response persisted throughout the first week, with RER oscillating between 0.8 and 1.0 across the 24-hour cycle, consistent with continued mixed-substrate oxidation and incomplete transition to sustained lipid utilization. The shift to a stabilized metabolic state emerged progressively after the first week. While AL animals maintained stable RER profiles across the entire four-week period, RER in CR animals began to diverge after the first week: daytime values fell below 0.8, signaling a shift toward whole-body lipolysis, and by the second week a new metabolic steady state had emerged (**Fig. 1b**).

Based on these observations, we designated Days 3 and 7 as the early adaptation period, Days 14–15 as the late adaptation period, and Day 28 as the established steady state. For subsequent analyses, Day 13 was used as a proxy for Day 14 due to a technical interruption in data acquisition. Overlaying the longitudinal RER profiles confirmed this temporal separation: Days 3 and 7 clustered distinctly from Days 13 and 28, with the second-week profile displaying the high-amplitude diurnal substrate switching characteristic of classical CR paradigms^37,38^ (**Fig. 1d**). AL RER curves, in contrast, were indistinguishable across all timepoints (**Fig. 1c**).

The two-week adaptation threshold was equally apparent in EE dynamics. Under baseline conditions and throughout the AL period, EE displayed a consistent bimodal pattern: a broad peak at dark-cycle onset and a secondary sharp peak immediately preceding the light cycle (**Fig. 1e**). In the early phase of CR, this bimodal structure was preserved but remodeled; the secondary pre-light peak was attenuated, while the EE spike at ZT12 was amplified relative to AL controls (**Fig. 1e**). By the second week, however, the EE profile had reorganized entirely. The pre-light peak disappeared, giving way to a single high-magnitude peak at ZT12 followed by a continuous, linear decline across the remainder of the 24-hour cycle, a pattern that remained stable through Day 28 (**Fig. 1f**). VO_2_ and VCO_2_ profiles mirrored these EE dynamics across all timepoints (**Supplementary Fig. 1**). Ambulatory activity and feeding behavior co-varied closely with EE, suggesting that diurnal fluctuations in whole-body energy expenditure are predominantly shaped by behavioral and feeding activity (**Extended Data Fig. 1**). Together, these data indicate that approximately 14 days are required for mice to fully restructure whole-body fuel utilization and energy expenditure in response to CR.

Gross body weight in CR animals exhibited pronounced diurnal oscillations that were detectable within the first 24 hours of restriction and became increasingly prominent through the second week (**Fig. 2a–d**). Because CR animals consumed their entire daily allotment in a single feeding bout (**Fig. 2e–g**), we asked how much of this fluctuation was attributable to gastric content rather than true changes in body mass. Stomach weights were measured at Days 3 and 28 across the feeding window. On Day 3, stomach weight in CR animals increased by approximately 1 g at ZT15 relative to AL controls, a modest effect consistent with incomplete consolidation of feeding behavior during early adaptation (**Fig. 2h**). By Day 28, stomach weight in CR animals equaled the full mass of the ingested food at ZT15 and remained elevated for up to 8 hours post-feeding (**Fig. 2i**). The diurnal body weight oscillations in Fig. 2b therefore largely reflect the physical mass of the consolidated meal rather than meaningful fluctuations in body composition, and true body weight in CR animals was likely stable across the four-week period.

**Figure 2.**
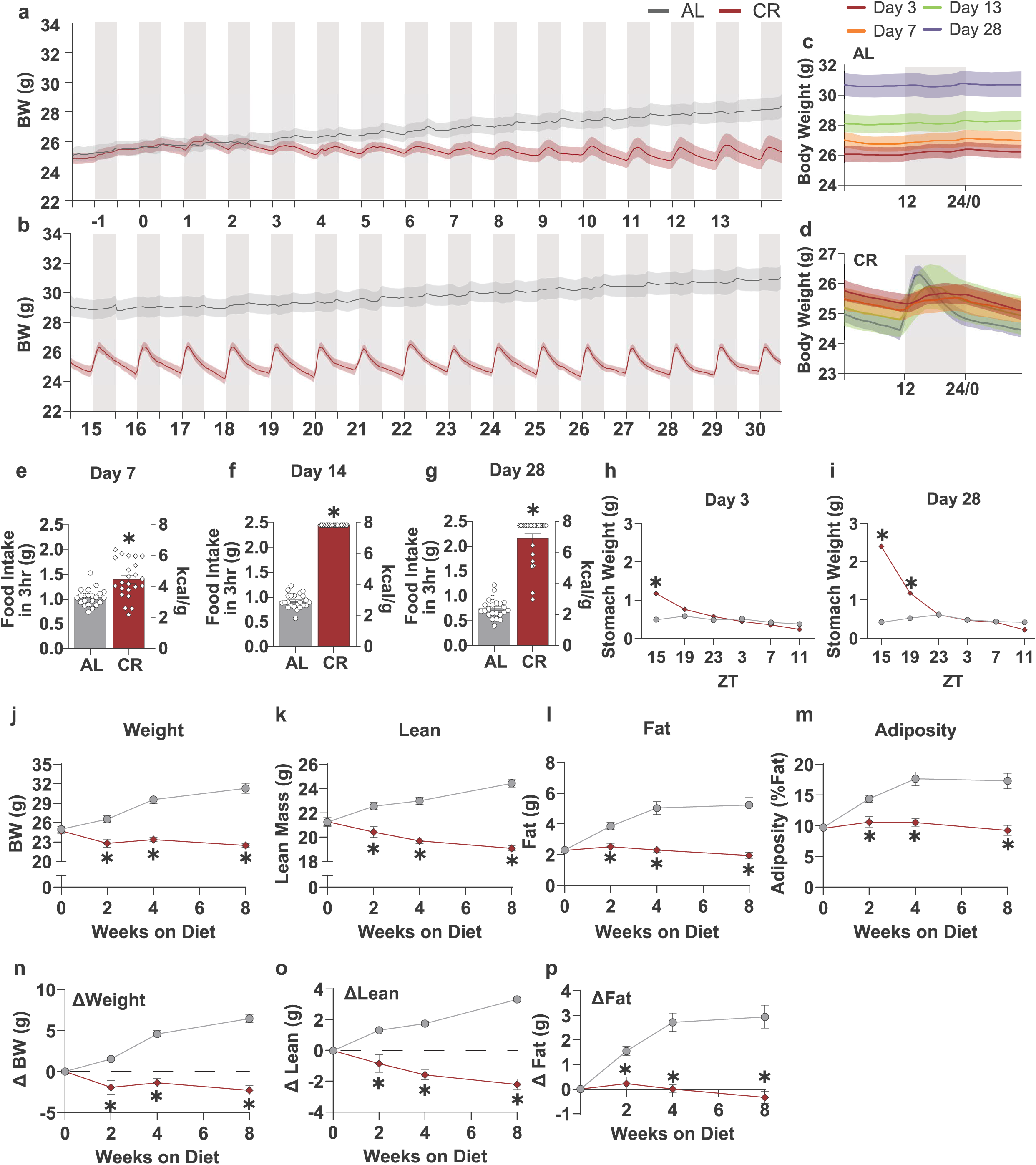
Body weight dynamics, feeding consolidation, and body composition during adaptation to CR. **a, b,** 4 week time course recording of body weight of AL and CR groups (n = 10-12). **c, d,** Overlay of longitudinal bodyweight profiles for AL and CR animals at Days 3, 7, 13, and 28 (n = 10-12). **e, f, g,** 3 h food intake measurement for AL and CR cohorts on days 7, 14 and 28 (n = 10-12). **h, i,** stomach weights at Day 3 and 28 at ZT 15, 19, 23, 3, 7 and 11 with ZT 15 representing the post-prandial state (fed from ZT 12-15) and ZT 11 representing the pre-prandial state (∼20 h fast) (n = 4 per timepoint). **j, k, I, m,** Longitudinal absolute body weight, lean mass, fat mass and adiposity measurement over eight weeks for AL and CR cohorts (n = 23-24) and **n, o, p** change in weight, lean mass and fat mass compared to baseline (Day 0). *P < 0.05 by two-way ANOVA with Sfdak’s multiple comparisons test.

Longitudinal body composition analysis over eight weeks confirmed this interpretation. CR animals maintained stable total body weight and fat mass from weeks 2 through 8, while lean mass declined progressively, consistent with prior reports (**Fig. 2j-p**)^36,37^. AL animals, by contrast, showed continuous accretion of total body weight, fat mass, and lean mass across the same period (**Fig. 2j-p**).

### Caloric restriction stabilizes circulating glucose and suppresses ketogenesis through consolidated feeding

To characterize the systemic metabolic response to CR across the adaptation timeline defined above, we measured circulating blood glucose and beta-hydroxybutyrate (βHB) at Days 7, 14, and 28. On each day, independent cohorts were subjected to 4-, 12-, or 20-hour fasting intervals beginning at ZT15, immediately following a 3-hour feeding window (ZT12–ZT15). Pre-feeding body weights were recorded at ZT12.

At Day 7, pre-feeding body weights were equivalent between AL and CR cohorts (**Fig. 3a**), and CR mice trended toward greater absolute food intake during the feeding window, though only the 12-hour fasting cohort reached statistical significance (**Fig. 3b**). Following the 4-hour fast, CR mice exhibited lower blood glucose than AL controls; notably, this concentration remained stable across the 12- and 20-hour intervals, whereas AL glucose peaked at 4 hours and declined progressively thereafter (**Fig. 3c)**. βHB concentrations did not differ significantly between groups at any fasting timepoint on Day 7 (**Fig. 3d**).

**Figure 3.**
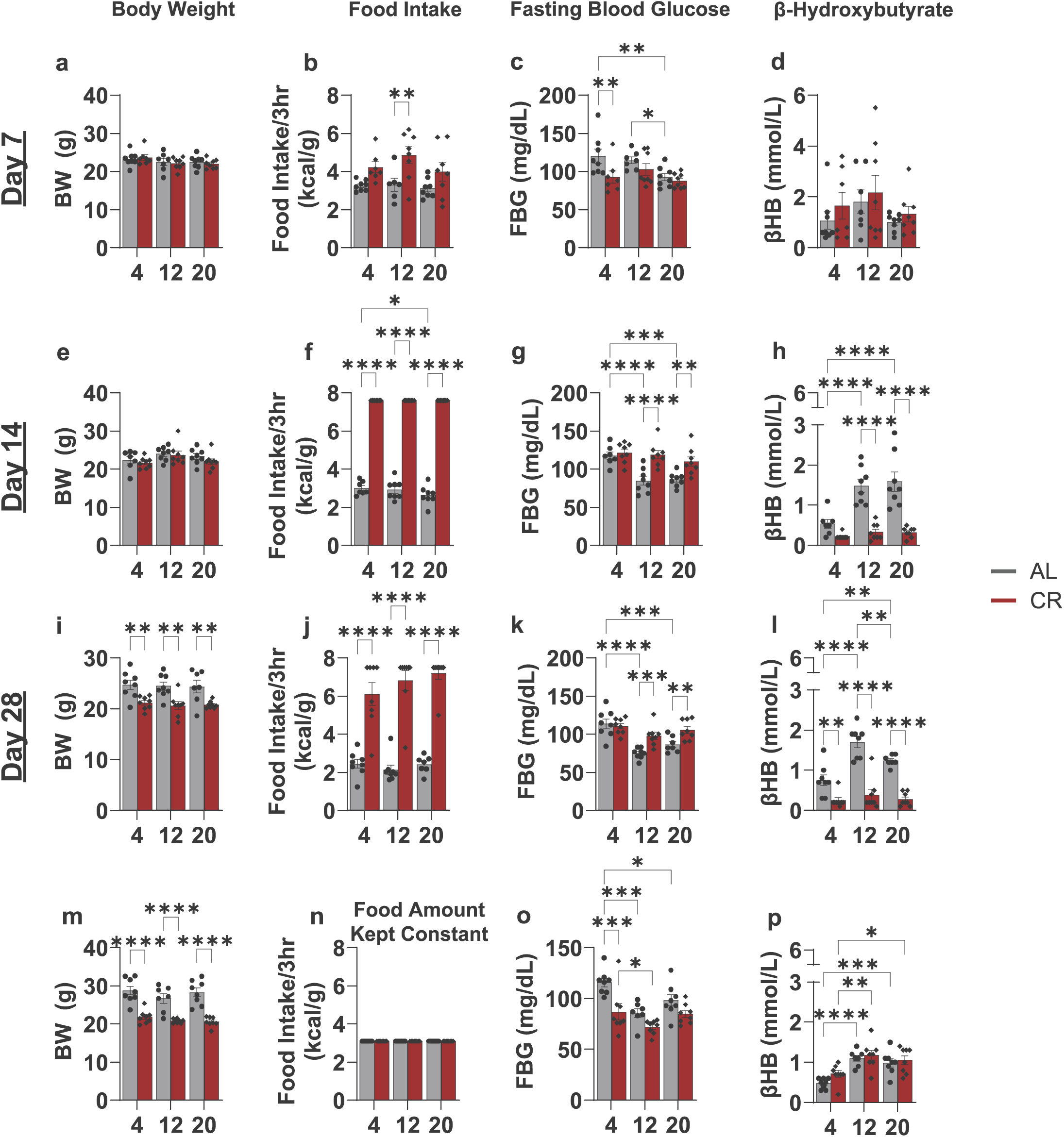
Caloric restriction stabilizes circulating glucose and suppresses ketogenesis through consolidated feeding. Independent cohorts of AL and CR mice were subjected to 4-, 12-, or 20-hour fasting intervals beginning at ZT15 following a 3-hour feeding window (ZT12-ZT15) on Days 7, 14, and 28 of the intervention. **a, e, i,** Pre-feeding body weights at ZT11 for AL and CR cohorts on Days 7, 14, and 28, respectively. **b, f, j,** Absolute food intake during the 3-hour feeding window for each fasting interval cohort (4, 12, or 20 hours) on Days 7, 14, and 28, respectively. **c, g, k,** Blood glucose concentrations at 4, 12, and 20 hours post-feeding for AL and CR cohorts on Days 7, 14, and 28, respectively. **d, h,** I, Circulating beta-hydroxybutyrate (f3HB) concentrations at 4, 12, and 20 hours post-feeding for AL and CR cohorts on Days 7, 14, and 28, respectively. **m,** Pre-feeding body weights at ZT11 for AL and CR cohorts under volume-matched feeding conditions. **n,** Absolute food intake during the 3-hour feeding window under volume-matched conditions (1 g chow). **o,** Blood glucose concentrations for AL and CR cohorts under volume-matched feeding conditions. **p,** Circulating f3HB concentrations for AL and CR cohorts under volume-matched feeding conditions. Data are mean± s.e.m.; n = 7-8 per group per timepoint. *P < 0.05 by two-way ANOVA with Sidak’s multiple comparisons test.

By Day 14, feeding behavior had markedly consolidated, CR animals consumed more than twice the absolute food mass of AL controls during the 3-hour window despite equivalent pre-feeding body weights (**Fig. 3e–f**). The glucose response had also shifted; CR animals now maintained stable blood glucose concentrations across all three fasting intervals, a plateau that contrasted sharply with the progressive decline observed in AL controls (**Fig. 3g)**. Most notably, while AL animals exhibited a fasting-duration-dependent rise in βHB, CR animals maintained near-basal βHB concentrations at every timepoint (**Fig. 3h**). By Day 28, pre-feeding body weights in CR animals had fallen significantly below those of AL controls; however, the patterns of feeding consolidation, glucose stability, and βHB suppression were maintained across all fasting intervals (**Fig. 3i–l**).

The persistent suppression of ketogenesis across extended fasting intervals was unexpected and warranted further investigation. Given that CR stomach mass remained elevated for up to 8 hours post-feeding (**Fig. 2i**), we reasoned that the physical volume of the consolidated meal may delay gastrointestinal transit, extending nutrient delivery well into the nominal fasting period. To test this, we restricted CR animals to a volume-matched ration equivalent to the typical 3-hour AL intake (**Fig. 3n**). Under these conditions, CR mice exhibited lower blood glucose than AL controls but produced circulating βHB concentrations equivalent to those of AL animals (**Fig. 3o–p**). These findings suggest that the large meal volume inherent to consolidated CR feeding prolongs gastric emptying and nutrient absorption, sustaining systemic nutrient availability during what is conventionally classified as the fasting period and thereby suppressing ketogenesis independently of fasting duration.

### Identification of transcriptional networks underlying the transition from ad libitum to caloric restriction

Having established that whole-body metabolic adaptation to CR progresses through distinct phases, with Days 3 and 7 capturing early adaptation, Day 14 late adaptation, and Day 28 the metabolic steady state, we next sought to characterize the hepatic transcriptional dynamics underlying this transition. Liver tissue was collected at four-hour intervals (ZT 15, 19, 23, 3, 7, and 11) across all four adaptive timepoints from both CR and AL animals and prepared for bulk RNA sequencing (**Fig. 4a**). Principal component analysis (PCA) of the resulting data revealed a progressive separation of CR transcriptional profiles from AL controls between Days 3 and 28, while AL profiles remained stable across the same period (**Supplementary Fig. 2b**). Time of day emerged as a primary source of variance driving cluster separation (**Supplementary Fig. 2c**), indicating that CR remodels the hepatic transcriptome in a manner shaped by both the duration of the intervention and the time of tissue collection.

**Figure 4.**
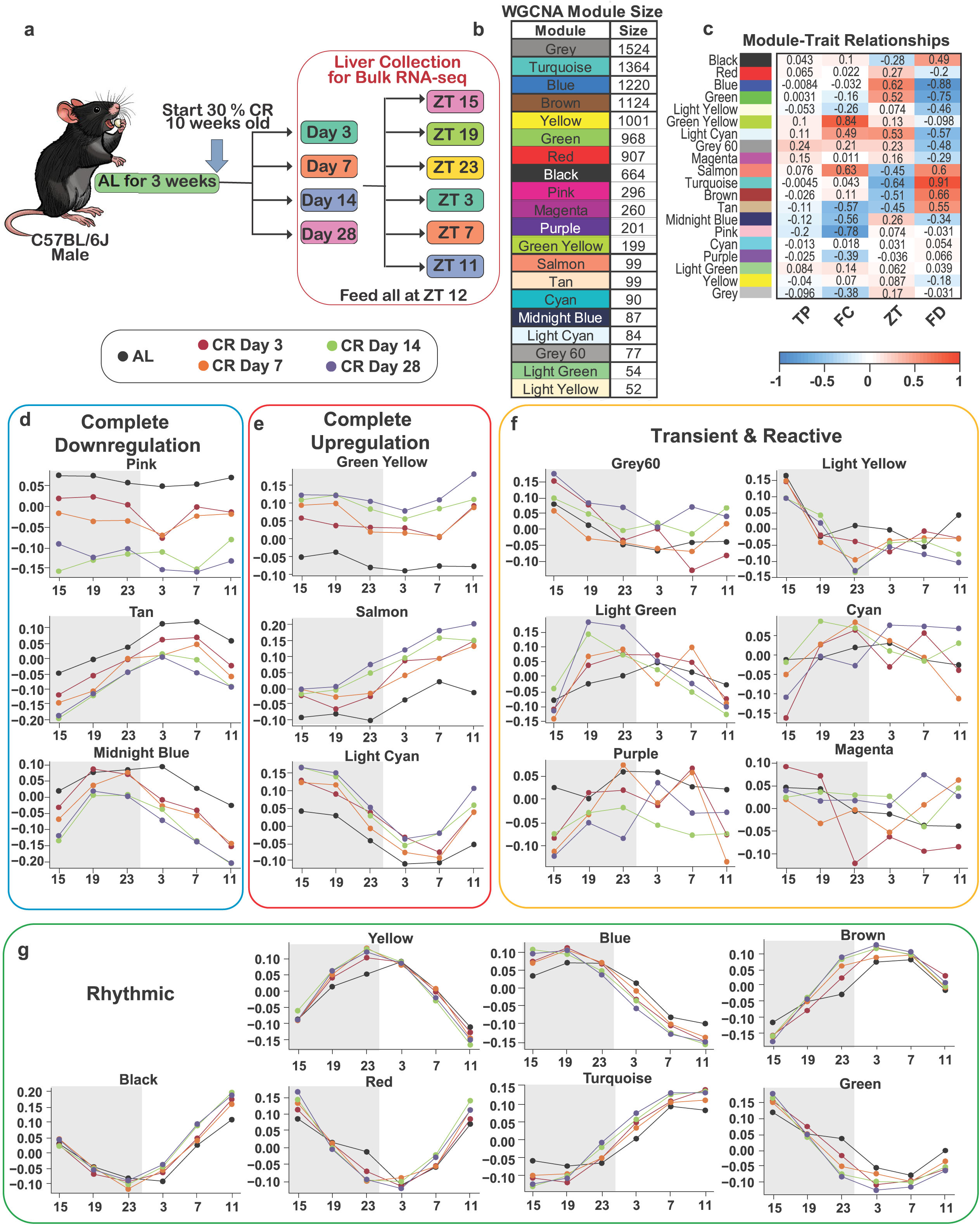
Hepatic WGCNA identifies 19 co-expression modules with distinct relationships to experimental variables. **a,** Experimental schematic illustrating the tissue collection design across four adaptive timepoints (Days 3, 7, 14, and 28) and six ZT timepoints (15, 19, 23, 3, 7, and 11) for bulk RNA sequencing. **b,** Module color assignments depicting the 19 non-overlapping co-expression modules identified by WGCNA of the liver transcriptome. **c,** Module-trait relationship heatmap depicting Pearson correlation coefficients between module eigengenes and experimental variables: transition period (TP), feeding condition (FC), time of day (ZT), and fasting duration (FD). Color scale represents correlation magnitude; asterisks denote statistical significance. **d-g,** Average module eigengene expression profiles stratified by ZT and adaptive timepoint, illustrating the classification of iWAT modules into **(d)** complete downregulation **(e)** complete upregulation, **(f)** transient/reactive, and **(g)** rhythmic functional clusters. Data are presented as average per group. n = 3 biological independent mice per feeding condition per adaptive timepoint per ZT timepoints.

While PCA captured the overarching trajectory of adaptation, it lacked the resolution to isolate discrete co-regulated gene networks underlying these global shifts. To systematically deconstruct this multidimensional variance, we applied weighted gene co-expression network analysis (WGCNA). Using a dynamic tree-cut algorithm, we identified 19 non-overlapping co-expression modules, each designated by a unique color (**Fig. 4b; Supplementary Fig. 3**), and assessed their correlation with four experimental variables: transition period (TP), diet, time of day (ZT), and approximate fasting duration (FD).

Fasting duration emerged as the dominant driver of transcriptional variation across the network. The Turquoise module exhibited the strongest positive correlation with FD (r = 0.91), reflecting progressive upregulation with increasing fasting time, while the Blue module showed a strong negative correlation (r = −0.88), consistent with transcriptional downregulation during the fasting window (**Fig. 4c**). The Green (r = −0.75) and Brown (r = 0.66) modules were also significantly associated with fasting duration. Beyond this variable, distinct modules segregated by dietary intervention: the Green Yellow module displayed the strongest positive correlation with CR (r = 0.84), the Salmon module showed a moderate positive correlation (r = 0.63), and the Pink module exhibited a strong negative correlation. In contrast, module correlations with TP were unexpectedly weak, the highest was observed in the Grey60 module (r = 0.24), despite TP driving clear cluster separation in PCA (**Supplementary Fig. 2c**).

We reasoned that the dominant variance attributed to ZT and FD was masking the contribution of TP to global network correlations. To decouple these variables, we stratified the data by ZT and TP, isolating module-specific eigengene values across both factors (**Fig. 4d**). This stratification enabled the classification of hepatic transcriptional networks into four functional categories: complete downregulation, complete upregulation, transient/reactive, and rhythmic. Where modules exhibited overlapping characteristics, for example, the Light Cyan module displayed both rhythmicity and a global shift in expression relative to AL controls, they were assigned to the upregulation or downregulation categories, as the absolute magnitude of expression at every ZT remained distinct from AL controls.

Three modules exhibited comprehensive transcriptional downregulation under CR. The Pink module showed the most pronounced effect, it lacked detectable rhythmicity and underwent an immediate, persistent downregulation apparent as early as Day 3 (**Fig. 4d**). The Tan and Midnight Blue modules followed a similar trajectory of progressive downregulation but retained rhythmic expression profiles throughout. Conversely, the Salmon, Light Cyan, and Green Yellow modules exhibited marked transcriptional upregulation under CR. Mirroring the downregulated group, this set comprised one non-rhythmic module (Green Yellow) and two with preserved rhythmicity (**Fig. 4e**). The Green Yellow module showed a pronounced increase in expression at ZT11, corresponding to the late pre-prandial state, suggesting that its constituent gene networks may be engaged in the food-anticipatory response. In contrast to the complete up- and downregulated groups, the rhythmic group, comprising the Yellow, Red, Turquoise, Brown, Black, Green, and Blue modules, showed minimal deviation from AL profiles. These modules maintained their characteristic waveforms across the transition, adapting primarily through amplitude scaling rather than phase shifts. The transient/reactive group exhibited a distinct profile characterized by acute, unstructured bursts of expression, likely reflecting an immediate transcriptional stress response during the early phase of adaptation to CR (**Fig. 4f**).

### Soft-thresholding and differential expression analysis resolve four distinct hepatic transcriptional adaptation patterns

To capture genes with pleiotropic regulatory roles, we expanded each module beyond strict cluster assignments using a soft-thresholding approach based on module membership scores (|kME| > 0.6). For example, applying this threshold to the Light Cyan module increased the gene set from 84 to 381 transcripts. This expanded framework was used to identify hub genes operating across multiple regulatory contexts and provided a more robust dataset for KEGG and Reactome pathway enrichment analyses. Differentially expressed genes (DEGs; adjusted P < 0.05) were then identified from the kME-defined datasets at each transition period (Days 3, 7, 14, and 28). UpSet plots were used to visualize the intersection of DEGs across the duration of the intervention, enabling identification of gene subsets that characterize the early adaptive response versus those defining sustained states (**Extended Data Fig. 2**). This analysis resolved four distinct patterns in the hepatic transcriptional profiles during adaption to CR.

The early transition & sustained cluster (Green Yellow, Pink, Light Cyan, and Salmon modules) captured the acute phase of hepatic transcriptional remodeling (**Extended Data Fig. 2a**). Genes within these modules were differentially expressed as early as Day 3, after which expression profiles either stabilized or continued to accumulate DEGs through Day 28. The Green Yellow module established a transcriptional state at Day 3 that remained largely static, retaining 85% of its initial gene pool (n = 116) through Day 28 without further increases. The Pink, Light Cyan, and Salmon modules, by contrast, showed continuous expansion following the initial transition. The Pink module increased from 64 to 195 DEGs while fully preserving the expression state established at Day 3; the Light Cyan and Salmon modules followed a similar trajectory, each comprising a distinct set of DEGs at Day 3 (n = 95 and n = 114, respectively) that more than doubled by Day 28 (n = 222 and n = 314, respectively).

The late transition & sustained cluster comprised the Tan, Blue, Midnight Blue, Red, and Light Yellow modules (**Extended Data Fig. 2b**). Rather than reaching maximal DEG counts during the acute phase, these modules initiated their response primarily at Day 7 and stabilized by Day 14. The Midnight Blue and Tan modules showed continuous increases from Day 3, stabilizing on Day 14. The Blue and Red modules displayed a non-monotonic profile, maintaining modest numbers of altered transcripts during the first week before a pronounced increase at Day 14 (71 to 197 and 73 to 182 DEGs, respectively), followed by stabilization. The Light Yellow module exhibited the most delayed response within this cluster, remaining largely unchanged at Days 3 and 7 (n = 9 and n = 36, respectively) before an approximately tenfold increase at Day 14 (n = 92) that was maintained through Day 28 (n = 96).

The progressive remodeling cluster (Turquoise, Brown, Black, Green, Yellow, and Light Green modules) was defined by continuous increase in DEG counts during the later adaptive timepoints (**Extended Data Fig. 3a**). These modules showed minimal divergence from AL controls during the acute phase, with significant number of transcriptional changes emerging only as CR advanced into the established phase. The Turquoise module increased from 66 DEGs on Day 3 to 300 by Day 28. The Brown module similarly accelerated late in the intervention, reaching 265 DEGs by Day 28, with 46% of constituent transcripts uniquely present at this final timepoint. The Black and Green modules maintained a latent profile through Day 7 (n = 68 and n = 77, respectively) before a sharp increase at Day 14 (n = 208 and n = 202, respectively). The Yellow and Light Green modules reflected more gradual trajectories: the Yellow module increased to 151 DEGs by Day 14 and 173 by Day 28, while the Light Green module expanded modestly from 14 DEGs at Day 3 to 36 by Day 28.

The late-onset cluster (Grey60, Cyan, Magenta, and Purple modules) was defined by a near-complete absence of DEGs during the acute transition (**Extended Data Fig. 3b**). Transcriptional profiles within these modules remained unchanged through Day 7, requiring two full weeks of CR before significant changes emerged. The Cyan module exhibited the most extreme delayed profile, with no detectable changes on Days 3 or 7 (n = 0) before a specific response at Day 14 (n = 28 DEGs). The Grey60 module maintained a near-baseline state at early timepoints (n = 6 and n = 7) before a discrete shift at Day 14 (n = 57). The Magenta and Purple modules followed a similar pattern, with minimal early divergence (n ≤ 4 DEGs) followed by approximately tenfold and thirteenfold increases, respectively, at Day 14 (n = 33 and n = 39 DEGs).

### Early kinless and connector hub genes define the scaffold of hepatic adaptation to caloric restriction

Having defined the temporal co-expression modules presiding over the hepatic transcriptional remodeling to CR, we next sought to identify the hub genes driving these coordinated signatures. To isolate upstream initiators, we applied a cartographic framework to genes exhibiting significant differential expression exclusively during the acute adaptive phase (Days 3 and 7), classifying each node by its within-module degree z-score (z ≥ 2.5) and inter-modular participation coefficient (P) to resolve positional roles within the global co-expression network (**Fig. 5a; Supplementary Fig. 6**)^39^. This approach delineated two tiers of pleiotropic influence among the early hub population.

**Figure 5.**
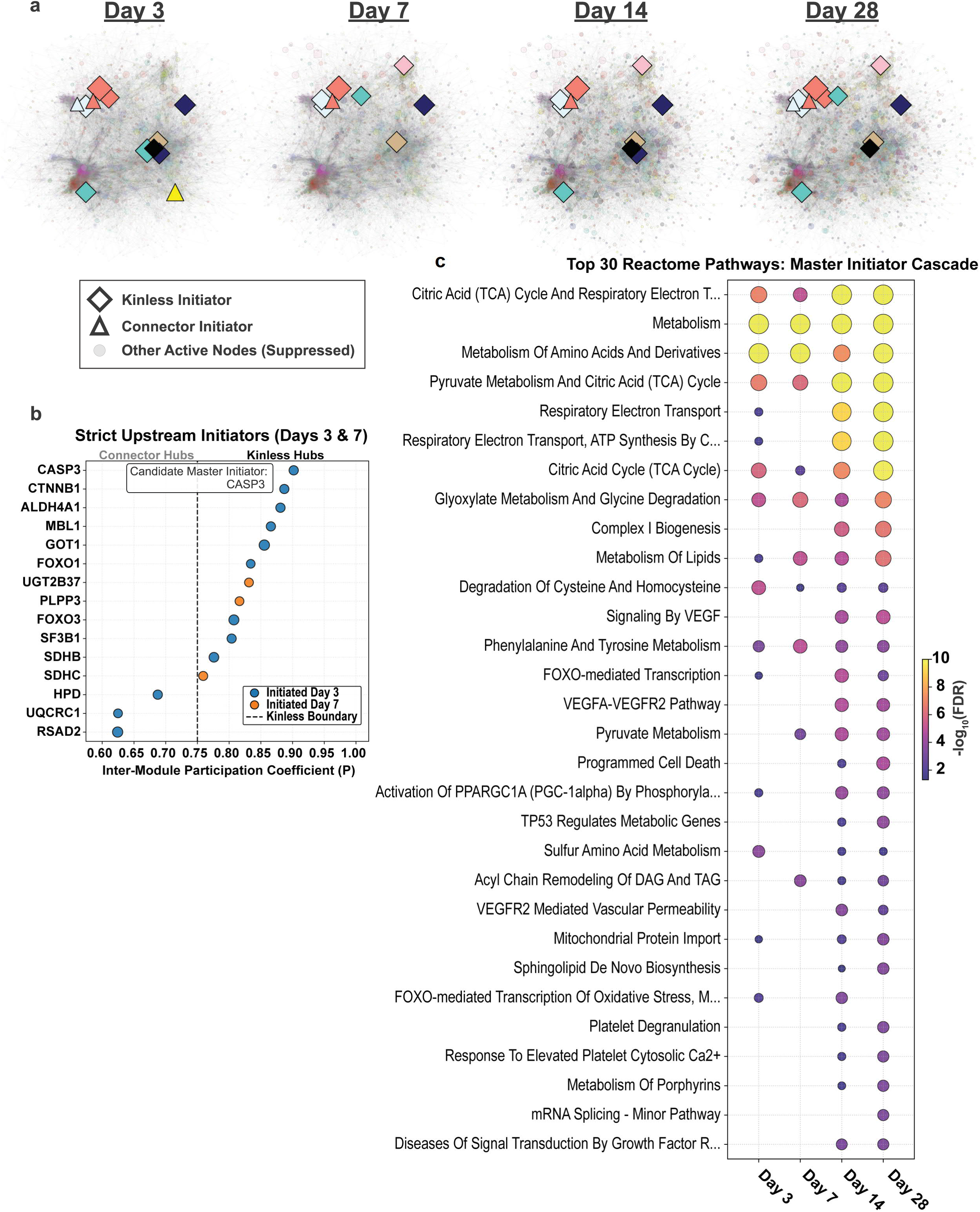
Early kinless and connector hub genes define the regulatory scaffold of hepatic adaptation to CR. **a,** Cartographic classification of genes with significant differential expression during the acute adaptive phase (Days 3 and 7), plotting within-module degree z-score (z ≥ 2.5) against inter-modular participation coefficient (P) to resolve positional roles within the global co-expression network. **b,** Identification of kinless hubs (P > 0.75) and connector hubs (0.30 < P :s; 0.75) among acute-phase DEGs. **c,** Toppe 30 reactome pathway enrichment analysis of the consolidated initiator network (hub genes and their first-degree topological neighbors). KEGG and GO enrichment analyses are shown in **Extended Data Fig. 4.**

Kinless hubs (P > 0.75), defined by connections distributed homogeneously across all modules, comprised *Ctnnb1, Aldh4a1, Mbl1, Got1, Foxo1, Ugt2b37, Plpp3, Foxo3, Sf3b1*, and *Sdhb* (**Fig. 5b**). Among these, Casp3 possessed the highest participation coefficient and was identified as the top candidate initiator of the hepatic CR response. A secondary tier of connector hubs (0.30 < P ≤ 0.75), including *Sdhc, Hpd, Uqcrc1*, and *Rsad2*, mediated more localized inter-modular communication, bridging adjacent functional clusters without the broad module-spanning connectivity that defines kinless nodes.

To define the functional consequences of these early hubs, the initiators and their first-degree topological neighbors were aggregated into a consolidated network and subjected to pathway enrichment analysis. Reactome analysis revealed pronounced enrichment of core metabolic and bioenergetic pathways (**Fig. 5c**). The most significantly enriched pathways, including the citric acid (TCA) cycle and respiratory electron transport, metabolism of amino acids and derivatives, and pyruvate metabolism, showed sustained activation that intensified through Days 14 and 28, consistent with the initiator network establishing a stable reprogrammed metabolic state rather than a transient stress response. The network also regulated discrete signaling circuits, including enrichment of VEGF signaling, FOXO-mediated transcription, and TP53-regulated metabolic gene programs which indicated that the kinless and connector hubs coordinate long-term transcriptional reprogramming in parallel with immediate bioenergetic remodeling (**Fig. 5c**).

These findings were also supported by KEGG and Gene Ontology (GO) enrichment analyses (**Extended Data Fig. 4**). KEGG enrichment confirmed regulatory involvement in steroid hormone biosynthesis, retinol metabolism, the citrate cycle, and thermogenesis (**Extended Data Fig. 4a**). GO Biological Process analysis identified mitochondrial ATP synthesis–coupled electron transport, the aerobic electron transport chain, and tricarboxylic acid metabolic processes as the primary functional outputs of the initiator network (**Extended Data Fig. 4b**). Collectively, these enrichment profiles indicate that this restricted subset of early kinless and connector hubs coordinates transcriptional remodeling spanning FOXO-mediated gene regulation, TP53-dependent metabolic targets, mitochondrial bioenergetic networks, and VEGF-associated microvascular remodeling, establishing a transcriptional profile that progressively consolidates into the adapted state observed by Day 28.

### CR-driven transcriptional remodeling of inguinal white adipose tissue is time-of-day dependent

Adipose tissue coordinates systemic energy homeostasis with the liver through lipid storage and mobilization^40–46^. To determine whether the transcriptional remodeling to CR in inguinal white adipose tissue (iWAT) parallels the remodeling observed in the liver, we performed bulk RNA-seq on iWAT collected across the same four adaptive timepoints and six circadian timepoints (**Fig. 6a**).

**Figure 6.**
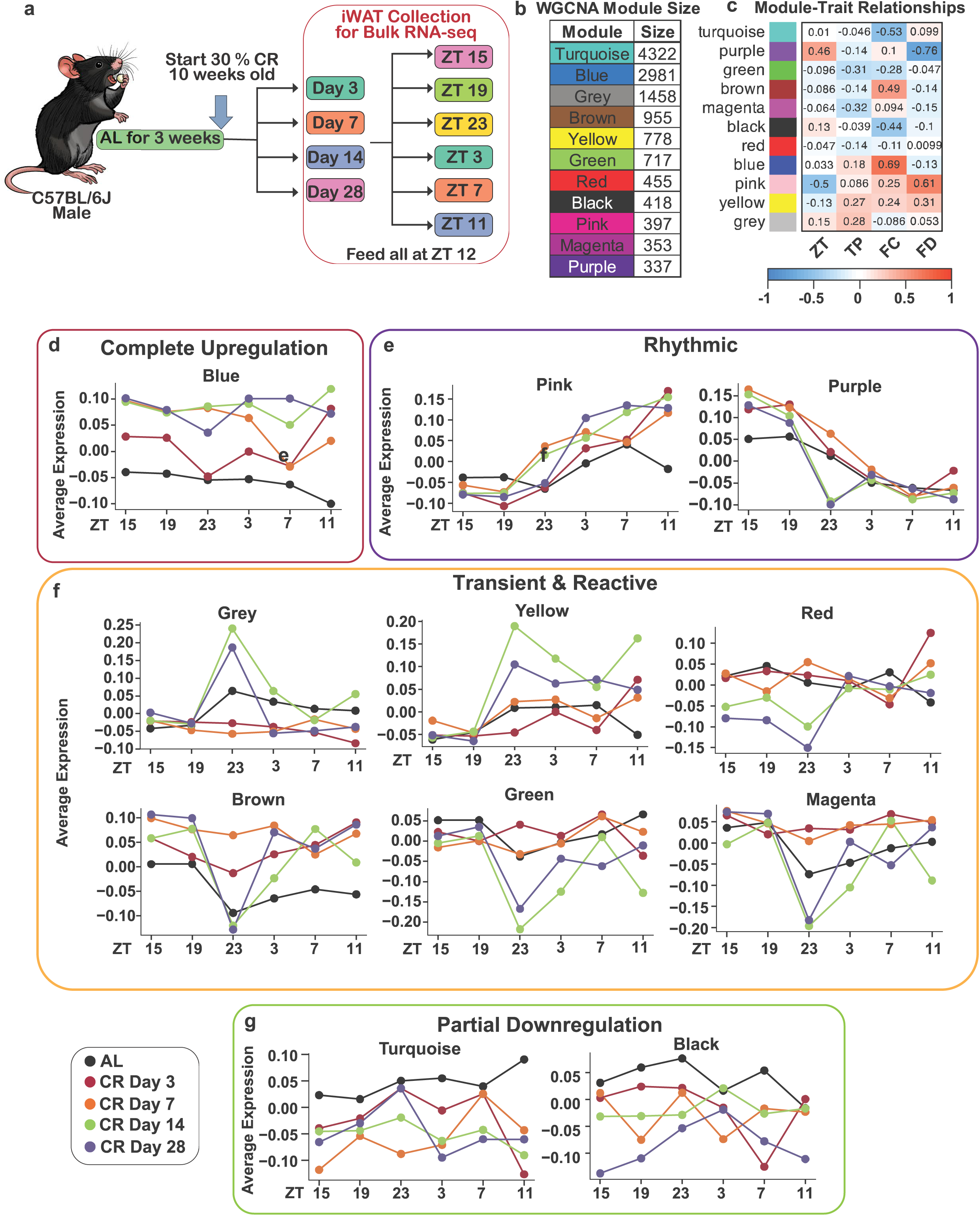
WGCNA of iWAT identifies time-of-day-dependent co-expression modules under CR. **a,** Experimental schematic illustrating the tissue collection design across four adaptive timepoints (Days 3, 7, 14, and 28) and six ZT timepoints (15, 19, 23, 3, 7, and 11) for bulk RNA sequencing. **b,** Module color assignments depicting the 11 non-overlapping co-expression modules identified by WGCNA of the iWAT transcriptome. **c,** Module-trait relationship heatmap depicting Pearson correlation coefficients between module eigengenes and experimental variables: transition period (TP), feeding condition (FC), time of day (ZT), and fasting duration (FD). Color scale represents correlation magnitude; asterisks denote statistical significance. **d-g,** Average module eigengene expression profiles stratified by ZT and adaptive timepoint, illustrating the classification of iWAT modules into **(d)** complete upregulation, **(e)** partial downregulation, **(f)** transienUreactive, and **(g)** rhythmic functional clusters. Data are presented as average per group. n = 3 biological independent mice per feeding condition per adaptive timepoint per ZT timepoints.

PCA of the iWAT dataset revealed a distinct pattern of variance compared to the liver. iWAT transcriptional profiles did not separate by dietary condition alone, nor did stratification by transition period resolve discrete clusters (**Supplementary Fig. 5a–b**). Instead, partitioning the dataset by time of day revealed divergence between AL and CR cohorts, with ZT23 exhibiting the strongest separation between dietary groups (**Supplementary Fig. 5c**). These findings indicate that iWAT transcriptional remodeling under CR is not uniformly distributed across the day but is concentrated at specific circadian timepoints.

### WGCNA identifies four functional clusters of iWAT co-expression modules under caloric restriction

WGCNA of the full iWAT dataset identified 11 non-overlapping co-expression modules (**Fig. 6b**). Module–trait relationship analysis revealed robust associations with specific experimental variables (**Fig. 6c**). The Blue module exhibited the strongest positive association with feeding condition (FC; r = 0.69), followed by the Brown module (r = 0.49). The Pink module was positively correlated with fasting duration (FD; r = 0.61), while the Purple module was positively associated with time of day (ZT; r = 0.46). In the opposing direction, the Purple module exhibited the strongest inverse correlation with FD (r = −0.76). The Turquoise and Black modules were inversely correlated with feeding conditions (r = −0.53 and r = −0.44, respectively), and the Pink module showed a negative association with ZT (r = −0.50). The Green and Magenta modules showed moderate negative correlations with transition period (TP; r = −0.31 and r = −0.32, respectively). In contrast to the hepatic dataset, where fasting duration emerged as the dominant driver of transcriptional variance, no single variable dominated the iWAT module-trait landscape, consistent with the PCA finding that iWAT transcriptional remodeling under CR is multifactorial and time-of-day dependent. Stratification of module eigengene expression across experimental timepoints organized these modules into four functional clusters: complete upregulation, partial downregulation, transient & reactive, and rhythmic (**Fig. 6d–g**).

The complete upregulation cluster comprised the Blue module (n = 2,981 transcripts), which exhibited a sustained increase in expression that emerged early in the intervention and remained elevated across subsequent adaptive timepoints (**Fig. 6d**). The partial downregulation cluster, containing the Turquoise (n = 4,322) and Black (n = 418) modules, displayed a progressive reduction in expression relative to AL controls, though this suppression was less pronounced than the corresponding downregulation observed in the liver (**Fig. 6g**).

The transient & reactive cluster comprised the largest proportion of transcriptomic variance, comprising the Grey (n = 1,458), Brown (n = 955), Yellow (n = 778), Green (n = 717), Red (n = 455), and Magenta (n = 353) modules (**Fig. 6f**). Rather than exhibiting stable directional changes, expression profiles within these modules were defined by interval-specific divergence from AL controls that was strictly dependent on time of day. The Grey, Brown, Yellow, and Magenta modules displayed coordinated expression shifts restricted to ZT23 and ZT3, consistent with the PCA findings (**Supplementary Fig. 5c**). The rhythmic cluster, comprising the Pink (n = 397) and Purple (n = 337) modules, maintained rhythmic expression profiles across both AL and CR conditions, suggesting that these gene networks preserve circadian regulation regardless of feeding condition (**Fig. 6e**).

These findings reveal that the iWAT transcriptome undergoes a temporally structured response to CR that is distinct from that of the liver. Whereas hepatic transcriptional remodeling was organized primarily by fasting duration and transition period, the iWAT response was predominantly shaped by time of day, with CR-driven divergence concentrated at specific circadian timepoints rather than distributed uniformly across the intervention.

### Module-specific DEG profiles reveal distinct clusters of iWAT adaptation to caloric restriction

As for the liver dataset, a soft-thresholding approach (|kME| > 0.6) was used to define module boundaries for the iWAT WGCNA modules, and DEGs (adjusted P < 0.05) were identified at each transition period (Days 3, 7, 14, and 28). UpSet plots were used to visualize DEG intersections across the intervention (**Extended Data Fig. 5**). This analysis resolved four adaptation clusters across the 11 modules: early-onset & constitutive, acute transient, bimodal, and late-onset.

The early-onset constitutive cluster (Blue and Turquoise modules) was characterized by substantial DEG counts at Day 3 and the most conserved transcriptomic profile across all adaptive timepoints (Extended Data Fig. 4a). The Blue module retained 90.9% of its initial Day 3 gene pool (n = 1,603 of 1,763) through Day 28. The Turquoise module similarly maintained 80.9% of its initial DEGs (n = 1,316 of 1,626) across all four timepoints. Both modules showed a transient increase in DEGs at Day 7, peaking at 2,223 and 2,911 total DEGs, respectively, before stabilizing at subsequent timepoints.

The acute transient cluster (Green, Pink, and Yellow modules) was defined by a sharp increase in DEG count at Day 7 (**Extended Data Fig. 5b**). The Green module expanded from 459 DEGs at Day 3 to 713 at Day 7, driven by a Day 7-specific set of transcripts that constituted 24.5% of the total Day 7 module volume and were not significant at any other timepoint. DEG counts within these modules subsequently attenuated by Day 14 and remained comparable to Day 3 levels through Day 28, consistent with a transcriptional signature restricted to the acute adaptive phase.

The bimodal cluster (Brown, Purple, Magenta, and Red modules) displayed a dual-peak pattern, recapitulating the initial Day 7 surge and subsequent Day 14 attenuation observed in the acute transient cluster before undergoing a secondary increase in DEGs at Day 28 (**Extended Data Fig. 5c**). The Brown module maintained a constitutive profile, retaining 86.6% of its Day 3 gene pool (n = 989 of 1,142), while its temporal fluctuations mirrored the non-monotonic behavior of the Purple, Magenta, and Red modules. This pattern was most pronounced in the Red module: following an initial increase to 204 DEGs at Day 7 and a decline to 146 at Day 14, a secondary expansion of 124 DEGs at Day 28, representing 48.2% of the terminal gene pool, established a late-stage transcriptional increase. The substantial Day 3 DEG counts in the Brown, Purple, and Magenta modules (up to n = 989) indicate that these networks share characteristics with the early-onset & constitutive cluster; their classification as bimodal is driven by the superimposed secondary recruitment event at Day 28 rather than by an absence of early activity.

The late-onset cluster comprised the Black module, which was largely unaffected by the Day 7 acute surge (**Extended Data Fig. 5d**). This module maintained a minimal transcriptional footprint at Day 3 (n = 28 DEGs) and showed progressive increases during the late adaptive stages, reaching 234 DEGs at Day 14 and 365 by Day 28. Of the terminal gene pool, 30.4% (n = 111 of 365) consisted of DEGs exclusive to Day 28, reflecting the recruitment of gene networks that emerge only after the acute response has subsided.

The iWAT DEG profiles reveal an adaptive profile distinct from that of the liver. Whereas hepatic transcriptional remodeling was organized as progressive, adaptive period-dependent restructuring, the iWAT response was dominated by an acute Day 7 surge followed by module-specific stabilization, resolution, or secondary recruitment, underscoring the tissue-specific nature of transcriptional adaptation to CR.

### Early hub architecture reveals a bifurcated transcriptional remodeling in iWAT adaptation to caloric restriction

To identify the topological hierarchy governing iWAT remodeling under CR, the cartographic framework applied to the liver was used to analyze the iWAT co-expression network (**Fig. 7a; Supplementary Fig. 7**). Given the time-of-day dependence of iWAT transcriptional remodeling identified by PCA, participation coefficients were calculated on day-averaged expression values to enable stable inter-modular connectivity estimates across the adaptive timeline. The candidate pool was restricted to genes with within-module degree z ≥ 2.5 and significant differential expression during the acute adaptive phases (Days 3 and 7), isolating a population of upstream hubs whose inter-modular connectivity was established prior to the broader transcriptional reorganization at later timepoints.

**Figure 7.**
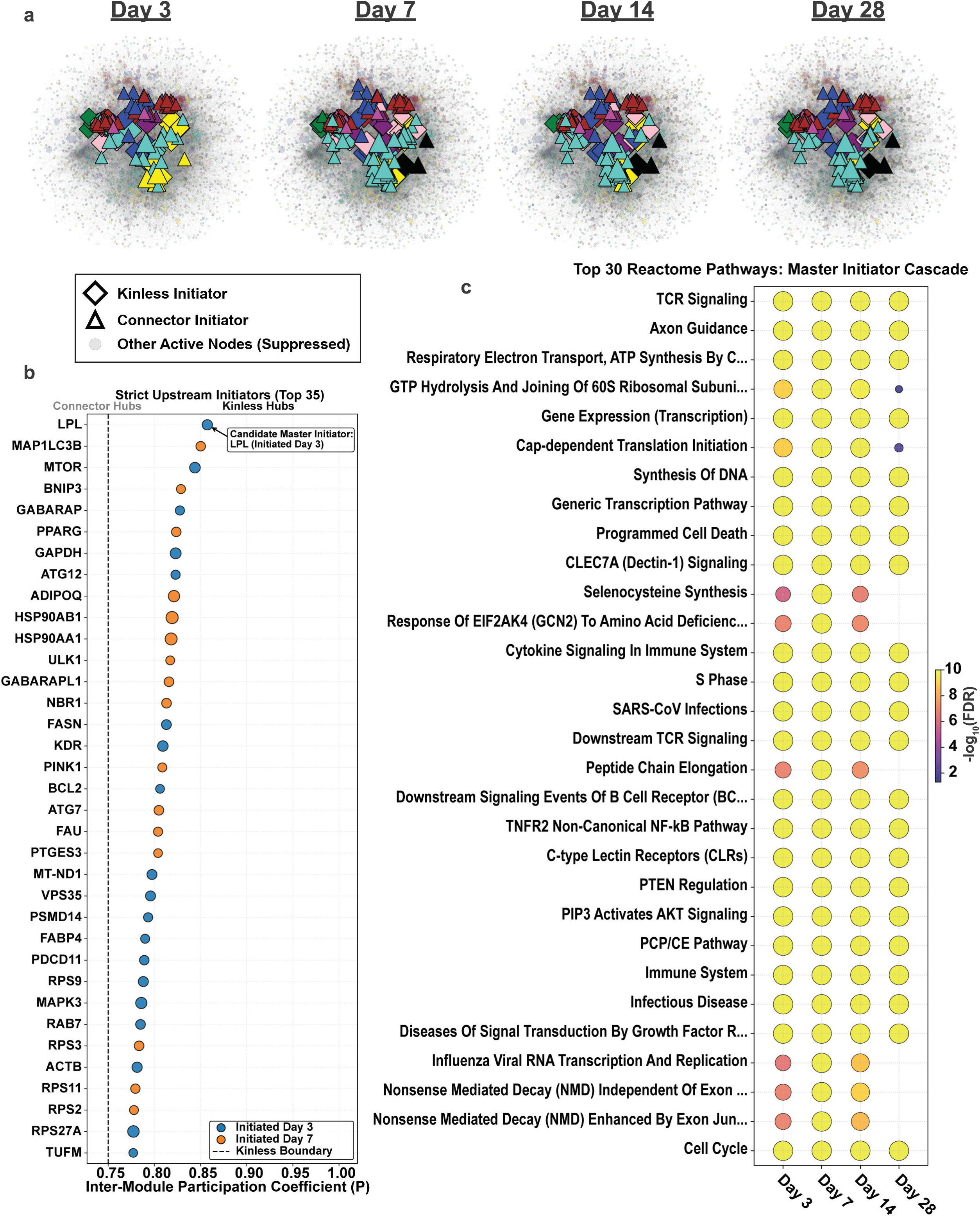
Early hub genes in iWAT reveal a bifurcated transcriptional profile during the adaption to CR. **a,** Cartographic classification of iWAT genes with significant differential expression during the acute adaptive phase (Days 3 and 7), using within-module degree z-score and participation coefficient. Participation coefficients were calculated on day-averaged expression values to account for time-of-day dependence. **b,** Identification of kinless and connector hubs, with Lpl exhibiting the highest participation coefficient. **c,** Tope 30 Reactome pathway enrichment analysis of the consolidated initiator network. KEGG and GO enrichment analyses are shown in **Extended Data Fig. 6**.

Within this hub population, *Lpl* emerged as the top candidate initiator, exhibiting the highest participation coefficient among kinless hubs (P > 0.75) across the network at Day 3. The broader kinless and connector hub population was enriched for regulators of adipocyte metabolism, macroautophagy, and nutrient sensing, including *Pparg, Adipoq, Fasn, Mtor, Map1lc3b, Ulk1, Pink1*, and *Atg7* (**Fig. 7b**). To characterize the functional output of these early hubs, the initiators and their first-degree neighbors were aggregated into a consolidated network and subjected to Reactome pathway analysis (**Fig. 7c**). This analysis revealed a bifurcated transcriptional profile. One arm was defined by enrichment of macromolecular biosynthetic and proliferative pathways, including GTP hydrolysis and joining of 60S ribosomal subunits, cap-dependent translation initiation, synthesis of DNA, and cell cycle progression. Concurrently, the same hub network showed enrichment of immunological and inflammatory pathways, involving TCR signaling, cytokine signaling in immune system, and CLEC7A (Dectin-1) signaling, a signature not observed in the hepatic initiator network.

These findings were supported by KEGG and Gene Ontology (GO) enrichment analyses (**Extended Data Fig. 6**). KEGG analysis confirmed enrichment of metabolic and catabolic programs, including oxidative phosphorylation, thermogenesis, and autophagy, alongside structural programs encompassing the ribosome and proteasome (**Extended Data Fig. 6a**). KEGG analysis further identified NF-κB, PI3K-Akt, and chemokine signaling as components of the immune-metabolic network coordinated by these hubs. GO Biological Process analysis identified ribosome biogenesis, rRNA metabolic processes, and cytoplasmic translation alongside cellular response to tumor necrosis factor, interleukin-1-mediated signaling, and stimulatory C-type lectin receptor signaling as functional outputs of the initiator network (**Extended Data Fig. 6c**).

These enrichment profiles suggest that the early kinless and connector hubs in iWAT coordinate a bifurcated transcriptional profile distinct from that of the liver. Rather than converging on a unified bioenergetic pathways, this network, anchored by *Lpl, Mtor*, and *Pparg*, directs concurrent activation of ribosomal biosynthetic and immune-regulatory programs, consistent with the simultaneous establishment of anabolic function and inflammatory response during early iWAT adaptation to CR.

### Cross-tissue transcriptional profiles during adaptation to caloric restriction identify a shared core transcript set

To identify transcriptional changes during adaptation to CR that were shared across tissues, DEGs from liver and iWAT (adjusted P < 0.05) were intersected across the four adaptive timepoints (Days 3, 7, 14, and 28), yielding a core set of 1,232 transcripts regulated in both tissues. Log2 fold-change (LFC) values for these shared transcripts were assembled into a matrix and subjected to unsupervised hierarchical clustering. A consensus eigengene, defined as the mean LFC across all module members at each timepoint, was computed for each resulting module (**Supplementary Fig. 8**). This approach separated the 1,232 transcripts into eight modules, visualized as a summary heatmap filtered to transcripts with |LFC| ≥ 1.0 (**Extended Data Figs. 7-8**).

Majority of modules displayed tissue-specific differences in the direction or magnitude of expression change despite being regulated in both tissues. Modules 1, 3, 4, 5, and 8 exhibited divergent expression trajectories between liver and iWAT (**Extended Data Fig. 7**): transcripts in Modules 1 and 3 were downregulated in iWAT and upregulated in liver, while those in Modules 4, 5, and 8 showed the opposite pattern. Modules 2 and 6 contained the largest number of DEGs and represented the two primary concordant responses: Module 2 was broadly downregulated in both tissues, and Module 6 was upregulated in both, though the magnitude of upregulation was greater in iWAT than in liver (**Extended Data Fig. 8**).

To assess whether the shared transcriptional response to CR intersected with genes implicated in longevity regulation, the 1,232 core transcripts were cross-referenced against the GenAge database. This identified 64 longevity-associated genes: 25 present exclusively in the human database, 27 exclusive to the model organism database, and 11 conserved across both. These genes were highlighted on the module heatmap (**Extended Data Figs. 7-8**); however, the majority fell below the |LFC| ≥ 1.0 visualization threshold at all adaptive timepoints examined.

Within the model organism database, distinct tissue-specific patterns were apparent at the level of individual genes. In Module 3, Mt2 was upregulated in liver and modestly downregulated in iWAT, with Fn1 showing a similar but attenuated pattern. Module 4 contained *Apoc3, Aco2*, and *Mpc1*, each of which was substantially upregulated in iWAT but showed only modest changes in liver. In Module 6, *Sod2, Cyc1, Sdhb*, and *Mdh2* were upregulated in iWAT with minimal corresponding changes in liver.

Among genes in the human longevity database, several deviated from the broader directional trend of their assigned module. In Module 2, *Pon1* was downregulated in iWAT but showed modest upregulation in liver, diverging from the concordant downregulation that defined the module. Module 8 contained *Mtor* and *Pparg*, both present in the human and model organism databases, as well as Gstp1 from the human database. *Mtor* was upregulated in iWAT and modestly downregulated in liver; *Pparg* and *Gstp1* were substantially downregulated in liver while remaining modestly upregulated in iWAT.

### Temporal dynamics of adipokine remodeling during adaptation to caloric restriction

Adiponectin and leptin are key adipose-derived signals linking tissue energy status to hepatic metabolic regulation, with receptors for both adipokines highly expressed in liver and iWAT. Given that the transcriptional remodeling observed in these tissues may be driven or reinforced by systemic endocrine signals, we asked whether circulating leptin and adiponectin profiles across the adaptive period were consistent with a coordinating role in the tissue-level gene expression changes identified by RNA-seq. Concentrations of both adipokines were quantified from plasma collected from the same animals used for RNA-seq, enabling direct alignment of endocrine and transcriptional data.

CR induced an acute post-prandial elevation in circulating leptin on Day 3, with significant increases at ZT15 and ZT23 relative to AL controls (**Fig. 8a**). This early elevation was partially maintained at Day 7, where leptin remained elevated at ZT15 but not ZT23 (**Fig. 8b**), indicating that the initial post-prandial leptin surge begins to resolve within the first week of restriction. By Day 14, leptin concentrations had decreased relative to AL controls (**Fig. 8c**). This decline was more pronounced by Day 28, where leptin remained significantly lower than AL controls from ZT23 through ZT11, consistent with progressive depletion of adipose leptin output over the course of CR (**Fig. 8d**).

**Figure 8.**
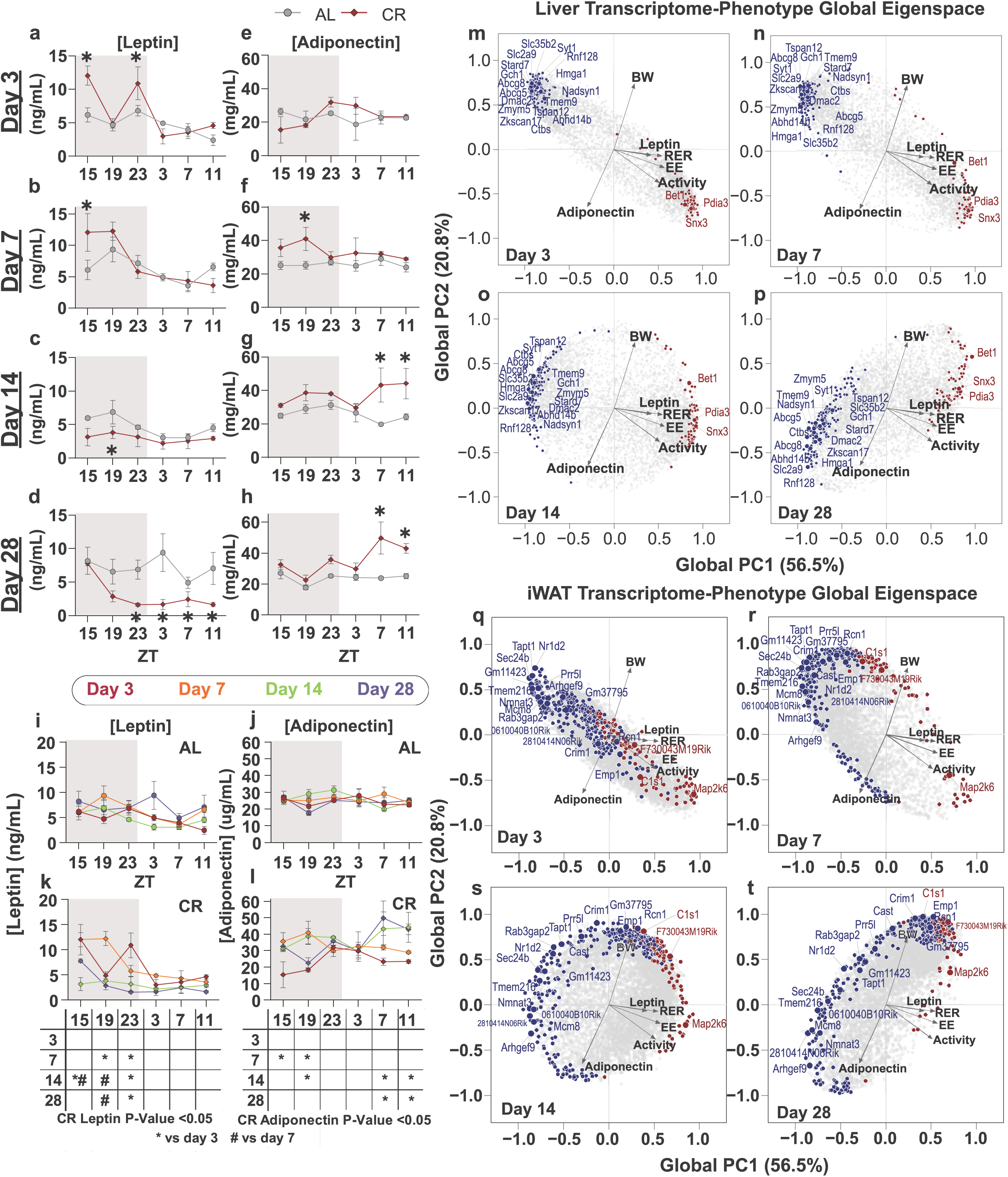
Adipokine dynamics and eigenspace projection reveal adiponectin-coupled transcriptional convergence during CR adaptation. **a-d,** Circulating leptin concentrations in AL and CR animals across six circadian timepoints at Days 3 **(a),** 7 **(b),** 14 **(c),** and 28 **(d). e-h,** Circulating adiponectin concentrations at Days 3 **(e),** 7 **(f), 14 (g),** and 28 **(h). i-j,** Within-group comparisons of leptin **(i)** and adiponectin U) across adaptive timepoints in AL animals. I, Inter-day comparisons of adiponectin in CR animals. **m-p,** Eigenspace projection of the hepatic RNA-seq dataset, with circulating leptin and adiponectin embedded as physiological reference vectors. Global PC1 (56.5% of variance) captures suppression of systemic metabolic output; Global PC2 (20.8%) resolves endocrine and somatic divergence. The top 5% of regulatory transcripts are tracked across Day 3 **(m),** Day 7 **(n),** Day 14 **(o),** and Day 28 **(p). q-t,** Eigenspace projection of the iWAT RNA-seq dataset across Day 3 **(q),** Day 7 **(r),** Day 14 **(s),** and Day 28 **(t).**

Adiponectin followed a distinct temporal pattern. On Day 3, CR animals did not differ from AL controls at any timepoint, indicating that adiponectin secretion is not acutely altered by the onset of CR (**Fig. 8e**). An increase was first detected on Day 7, restricted to ZT19 (**Fig. 8f**), consistent with an initial rise in adiponectin as adipose tissue begins to remodel under sustained negative energy balance. As CR progressed into the established adaptive phases, this elevation broadened and shifted later in the feeding–fasting cycle: by Days 14 and 28, significant increases were concentrated at ZT7 and ZT11, timepoints immediately preceding meal delivery, indicating that adiponectin secretion becomes preferentially elevated during the extended fasted state as CR is established (**Fig. 8g–h**).

Within-group comparisons confirmed the stability of AL profiles across all timepoints, with neither leptin nor adiponectin exhibiting significant inter-day variation over the 28-day period, providing a stable reference against which CR-induced changes could be assessed (**Fig. 8i–j**). In CR animals, inter-day comparisons revealed that leptin concentrations were most divergent during the early adaptive period: the post-prandial elevations present on Day 3 had largely resolved by Day 7, and the continued decline through Days 14 and 28 was most pronounced at timepoints distal from feeding (**Fig. 8j**). For adiponectin, inter-day divergence was concentrated at ZT15 and ZT19 between Days 3 and 7, consistent with the emergence of post-prandial elevation across the first week (**Fig. 8l**). By Days 14 and 28, divergence shifted to the pre-prandial window at ZT7 and ZT11, reflecting the progressive shift toward the extended fasted state as CR became established. These data indicate that leptin and adiponectin differ not only in the direction and magnitude of their changes under CR but also in the timing at which those changes become established – leptin declines rapidly and continuously from Day 3, whereas adiponectin undergoes a delayed increase that shifts toward the pre-prandial window only after the metabolic steady state is reached in the second week of restriction.

### Eigenspace projection reveals tissue-specific reorganization of regulatory networks across CR adaptation

The temporal dissociation between leptin and adiponectin suggests that these adipokines convey distinct endocrine information to their shared target tissues during CR adaptation. Their divergent trajectories map onto the same adaptive window in which the most pronounced transcriptional remodeling was identified, raising the possibility that these signals act in concert to coordinate the progressive reorganization of hepatic and adipose gene expression networks. To examine this relationship directly, liver and iWAT RNA-seq datasets were projected onto a shared phenotypic eigenspace in which circulating leptin and adiponectin were embedded as physiological reference vectors, enabling the temporal covariance between endocrine signaling and transcriptional remodeling to be quantified as spatial volatility (Δ) across the 28-day intervention (**Fig. 8m–t**). Global PC1 (56.5% of variance) captured the dominant suppresion of systemic metabolic output, with energy expenditure, RER, locomotor activity, and circulating leptin loading into the positive hemisphere, while Global PC2 (20.8%) resolved secondary endocrine and somatic divergence, establishing a structural opposition between body weight and circulating adiponectin. This coordinate system provided a stable physiological reference against which the spatial migration of the top 5% of regulatory transcripts was tracked across the intervention.

In the hepatic transcriptome, CR imposed an immediate and tightly clustered spatial bifurcation at Day 3. Baseline metabolic activators (including *Pdia3, Snx3*, and *Bet1*) aggregated in the lower-right quadrant, aligning with the energy expenditure and locomotor activity vectors, while CR-induced suppressive transcripts coalesced in the upper-left quadrant (**Fig. 8m**). Top regulatory drivers, including *Slc35b2* and *Syt1*, were diametrically opposed to the primary metabolic vectors and showed no structural correlation with the adiponectin axis, establishing the baseline topology for longitudinal tracking. Across Days 7 and 14, this initial polarization progressively attenuated as both networks dispersed outward, reflecting temporal rewiring of the hepatic regulatory landscape (**Fig. 8n–o**). By Day 28, the topography reached maximum spatial divergence, with *Rnf128* and *Syt1* exhibiting the greatest Euclidean displacement within the CR-induced network (**Fig. 8p**). *Rnf128* detached from its Day 3 coordinates and shifted into the lower-left quadrant to achieve terminal alignment with the adiponectin vector, indicating that the chronic hepatic CR state is characterized by a distinct adiponectin-coupled transcriptional program rather than sustained acute-phase signaling. Within the AL-associated network, volatile activators such as Bet1 tracked along the positive PC2 axis toward the body weight vector, consistent with somatic equilibration under prolonged CR.

An analogous but spatially distinct reorganization was observed in iWAT. At Day 3, AL-associated metabolic activators (including *Map2k6, C1s1*, and *F730043M19Rik*) aggregated in the lower-right quadrant in close alignment with the energy expenditure and locomotor activity vectors, while CR-induced suppressive transcripts (*Tapt1, Nr1d2, Sec24b*) coalesced in the left hemisphere without correlation to the adiponectin axis (**Fig. 8q**). Across Days 7 and 14, both networks diverged from their initial coordinates, the AL-associated network shifted along the positive PC2 axis toward the body weight vector, while the CR-induced network elongated across the negative PC1 hemisphere (**Fig. 8r–s**). By Day 28, the iWAT regulatory topography reached maximum divergence, with the AL-associated network consolidating in the upper-right quadrant (**Fig. 8t**). Volatile activators (*C1s1, F730043M19Rik, Gm37795*) aligned closely with the body weight vector, reflecting functional decoupling from primary metabolic outputs. Simultaneously, the CR-induced network exhibited a structural bifurcation: a cluster comprising *Crim1, Cast, Emp1*, and *Prr5l* remained anchored in the upper-left quadrant, whereas a volatile subset (including *Arhgef9, Nmnat3*, and *0610040B10Rik*) shifted inferiorly to align with the adiponectin vector. As in the liver, this terminal repositioning identifies an adiponectin-coupled transcriptional program as a feature of the stabilized CR state, suggesting that convergent endocrine coupling to adiponectin may represent a shared mechanism coordinating chronic adaptation across metabolically distinct tissues.

## Discussion

The transcriptional events that initiate and sustain adaptation to caloric restriction remain poorly defined, in part because most studies have characterized the adapted steady state rather than the transition itself ^29–34^. The redundancy of the signaling networks engaged by CR further complicates identification of individual regulatory nodes^15,21–28,47–51^. Here, we used continuous indirect calorimetry alongside longitudinal bulk RNA-seq of liver and iWAT across six circadian timepoints and four stages of adaptation to ask what changes first during the transition to 30% CR. We show that the two tissues organize their transcriptional adaptation through distinct temporal mechanisms, that restricted populations of early hub genes with high inter-modular connectivity precede the broader transcriptional remodeling in both tissues, and that dynamically regulated transcripts in both liver and iWAT converge on a shared adiponectin-coupled transcriptional state as CR is maintained.

Continuous metabolic monitoring revealed that the transition to CR proceeds through two discrete adaptive phases separated by a threshold at approximately 14 days. Prior studies have shown that CR-induced changes in body composition stabilize within the first two weeks^37,52^. Our calorimetry data suggest that the diurnal patterning of fuel utilization undergoes a parallel reorganization on the same timescale, with the transition to single-peak EE and sustained fat oxidation during the light cycle complete by Day 14. Characterization of the feeding-fasting dynamics within these adaptive phases revealed that circulating βHB remained near-basal even after 20-hour fasts at Days 14 and 28. The volume-matching experiment attributed this to prolonged gastric emptying from the consolidated bolus meal, indicating that the nominal fasting period in standard CR paradigms does not represent a true metabolic fast. This has implications for the broader field: tissue sampled during the light cycle of CR paradigms may reflect a fed rather than fasted metabolic state, and transcriptional changes attributed to fasting may instead arise from altered meal patterning, a distinction relevant to ongoing debates about the relative contributions of caloric deficit and time-restricted feeding^30,37,53–55^.

Hepatic transcriptional variance was driven primarily by fasting duration, with 19 co-expression modules dividing into four temporal DEG patterns defined by when separation from AL controls was first detected and how it accumulated. This transition period-dependent remodeling is consistent with evidence that the hepatic transcriptome is acutely sensitive to fasting duration^56–59^. In iWAT, no single variable organized transcriptional variance; separation between AL and CR profiles appeared only at specific timepoints, with ZT 23 showing the strongest divergence. The temporal DEG profile was also distinct from the liver, dominated by an acute Day 7 surge rather than progressive accumulation across the adaptation period. This time-of-day dependence of iWAT transcriptional changes under CR has not, to our knowledge, been previously described, and contrasts with the fasting duration-dependent pattern in the liver. Whether it reflects true circadian regulation or feeding-driven metabolic rhythms imposed by the consolidated meal remains to be determined. Together, these data suggest that the metabolic benefits of CR may arise from tissue-autonomous modes of transcriptional reorganization operating under different temporal regulation, rather than a single signaling cascade propagated across tissues. Consistent with this, the transcriptional boundaries in both tissues aligned with the metabolic phasing identified by calorimetry: hepatic DEG accumulation shifted most between Days 7 and 14, while the iWAT surge corresponded to the same metabolic transition.

Cartographic analysis of both co-expression networks identified small populations of kinless and connector hubs with differential expression restricted to the acute phase (Days 3 and 7) and inter-modular connectivity that preceded the broader remodeling at Days 14 and 28. In the liver, Casp3 had the highest participation coefficient, and the hub network was enriched for TCA cycle and respiratory electron transport, amino acid metabolism, FOXO-mediated transcription, and TP53-regulated metabolic genes. These enrichments increased at later timepoints, consistent with the acute-phase network initiating a metabolic state that is consolidated as adaptation proceeds. *Casp3* has been implicated in metabolic regulation independent of its apoptotic function^60–63^, and its identification as an early hub suggests a role in the initial transcriptional response to CR that warrants further investigation. In iWAT, *Lpl* had the highest participation coefficient, and the hub population included *Pparg, Adipoq, Fasn, Mtor, Map1lc3b, Ulk1, Atg7*, and *Pink1*. Unlike the liver, the iWAT hub network was functionally bifurcated, one arm enriched for ribosomal biosynthesis and cell cycle progression, the other for immune-regulatory pathways including TCR, cytokine, and CLEC7A (Dectin-1) signaling. This bifurcation is consistent with reports that adipose tissue adaptation to altered energy balance involves concurrent anabolic and immune-remodeling processes^45,64–66^, and the identification of *Pparg* and *Mtor* as early hubs rather than downstream effectors raises the possibility that these pathways function as initiators of adipose adaptation to CR.

Intersection of tissue-level DEGs identified 1,232 transcripts regulated in both liver and iWAT, including *Pparg* and *Mtor*, both present in the human and model organism GenAge longevity databases^67^. The majority of shared transcripts differed between tissues in the direction or magnitude of expression change, and longevity-associated genes showed the same pattern: *Sod2, Sdhb*, and *Mdh2* were upregulated in iWAT with minimal hepatic changes, while Pon1 was downregulated in iWAT but modestly upregulated in liver. These observations are consistent with emerging evidence that CR exerts tissue-dependent effects on the transcriptome^68,69^, and suggest that the relationship between CR and lifespan extension operates through tissue-specific modulation of shared regulatory targets.

The temporal profiles of circulating leptin and adiponectin provided a candidate mechanism for coordinating these tissue-autonomous transcriptional changes. Leptin declined rapidly from Day 3, while adiponectin increased with a delay, shifting to the pre-prandial window by Days 14 and 28, coinciding with the metabolic transition at which both tissues underwent their most substantial transcriptional remodeling. Projection of both RNA-seq datasets onto a shared phenotypic eigenspace revealed that dynamically regulated transcripts in both tissues shifted progressively across the adaptation period, and by Day 28 a subset in each, *Rnf128* in liver*, Arhgef9* and *Nmnat3* in iWAT, had aligned with the adiponectin vector. Adiponectin has well-characterized roles in hepatic fatty acid oxidation, insulin sensitization, and anti-inflammatory signaling^70–73^, and its identification here as a convergence point for tissue-specific transcriptional remodeling suggests it may function as an integrative endocrine signal linking acute metabolic perturbation to the chronic adapted state. Whether this coupling is causal will require interventional approaches such as adiponectin receptor blockade during the transition period.

This study has limitations. It was performed in male C57BL/6J mice, and generalizability to female animals, other strains, or other species is unknown. Bulk RNA-seq does not resolve cell-type-specific contributions, which is relevant for iWAT given the bifurcated enrichment of immune and metabolic pathways. High inter-modular connectivity during the acute phase does not demonstrate that a given hub gene is necessary or sufficient to initiate downstream adaptation; functional validation through targeted perturbation will be required to distinguish regulatory drivers from correlated nodes.

Our study identified small populations of early hub genes in both liver and iWAT whose inter-modular connectivity preceded and persisted through the broader transcriptional remodeling of CR adaptation. These hubs operated under distinct temporal regulation in each tissue yet converged on a shared adiponectin-coupled transcriptional state, providing candidate regulatory nodes for investigation of how the initial perturbation of caloric restriction is transduced into the sustained transcriptional changes that underlie its metabolic and longevity benefits.

## Materials and Methods

### Animals and feeding regimens

All animal procedures were approved by the Institutional Animal Care and Use Committee (IACUC) of the University of Texas Southwestern Medical Center and conducted in accordance with the National Institutes of Health Guide for the Care and Use of Laboratory Animals. Male C57BL/6J mice (age 7 weeks) were purchased from Jackson Laboratory and were placed on 2018 Teklad Global 18% Protein Rodent Diet acclimatized for ∼3 weeks before entering studies. All animals were housed individually under a 12:12 h light-dark cycled maintained at 23 ± 2°C. At 10 weeks of age, mice were randomized to either AL, ad libitum diet or CR, animals in which calories were restricted by 30%, and animals were fed once per day during the start of the dark period. Food was weighed and provided daily for CR group.

### Indirect Calorimetry and Metabolic phenotyping

Whole-body metabolic parameters were assessed by continuous indirect calorimetry using a Promethion metabolic cage system (Sable Systems International, Las Vegas, NV, USA) over a four-week period. Mice were acclimated to the metabolic chambers under ad libitum conditions for 48 hours prior to CR initiation to establish baseline metabolic profiles. Following acclimation, CR was initiated at ZT12. Oxygen consumption (VO_2_), carbon dioxide production (VCO_2_), respiratory exchange ratio (RER), energy expenditure (EE), locomotor activity, and food intake were recorded continuously at 5 minute intervals throughout the four-week recording period. Comprehensive metabolic data were recorded continuously over a 4-week period and analyzed in two distinct increments: Weeks 0–2 and Weeks 2–4. Locomotor activity was measured by an XYZ beam-break system. Data were analyzed using Macro Interpreter software (Sable Systems International) and visualized in GraphPad Prism (version 11). Mouse body composition was determined using an EchoMRI Body Composition Analyzer.

### Blood Glucose, Beta-Hydroxybutyrate, Adiponectin, and Leptin Measurements

To measure blood glucose and beta-hydroxybutryate levels across the adaptive timeline, independent cohorts of AL and CR mice were subjected to 4-, 12-, or 20-hour fasting intervals, starting the fast at ZT15 immediately following the 3-hour feeding window (ZT12–ZT15) on Days 7, 14, and 28 of the intervention. Pre-feeding body weights were recorded at ZT11. At the end of each fasting interval, tail vein blood was collected for measurement of blood glucose and circulating beta-hydroxybutyrate (βHB). Blood glucose was measured using a handheld glucometer (Contour Next), and βHB was measured using a handheld ketone meter (KetoBM). For the volume-matched feeding experiment, AL and CR animals was provided a restricted ration of 1 g chow (equivalent to the typical 3-hour AL intake) at ZT12 on Day 42 of CR. Blood glucose and βHB were measured at 4, 12, and 20 hours post-feeding as described above. After the collection of blood samples, adipokine and levels levels were assessed with Crystal Chem Mouse Adiponectin and Leptin ELISA Kit.

### Tissue Collection for RNA Sequencing

All animals were fed at ZT 12 and independent cohorts of AL and CR mice at six circadian timepoints (ZT15, ZT19, ZT23, ZT3, ZT7, and ZT11) across four adaptive timepoints (Days 3, 7, 14, and 28) of the CR intervention. Following blood collection via submandibular bleeding, mice were euthanized by rapid cervical dislocation, and tissues (liver and iWAT) were immediately, snap-frozen in liquid nitrogen, and stored at −80°C until processing. For Days 3 and 28, gross stomach weight was weighed.

### RNA Extraction and Bulk RNA Sequencing

RNA was extracted from liver or iWAT using TRI Reagent according to the manufacturer’s protocol (Sigma-Aldrich). The concentration and purity of RNA were determined by absorbance at 260/280nm using Nanodrop and RNA integrity was assessed using an Agilent TapeStation (Agilent Technologies); samples with an RNA integrity number (RIN) ≥ 7.0 were used for library preparation and sent to Novogene for sequencing.

### Read Quantification and Quality Control

Raw sequencing reads were quantified using Salmon (v1.10) with the --gcBias and --validateMappings flags against a mouse transcriptome index (mm39). Quality control was performed using FastQC (v0.11.8) and aggregated using MultiQC (v1.30).

### Transcript Quantification and Preprocessing

Transcript-level quantifications were imported into R using the tximeta package from Salmon output files. Gene-level counts were summarized using summarizeToGene() and a DESeq2 dataset object was constructed using diet, duration and ZT as covariates. Genes with fewer than 10 counts in fewer than 18 samples were excluded prior to downstream analysis.

### Differential Gene Expression Analysis

Differentially expressed genes were identified using DESeq2, comparing CR and AL conditions on filtered gene-level counts imported from Salmon. Ensembl gene identifiers were converted to gene symbols using the biomaRt package (v2.62.1). Log_2_ fold changes and adjusted P values (Benjamini–Hochberg correction) were used to construct volcano plots in ggplot2, with fold-change thresholds of ±1 and an FDR cutoff of 0.05. Genes meeting both thresholds were highlighted and labeled accordingly.

### Principal Component Analysis

Principal component analysis was carried out on count data that were transformed using the Variance Stabilixing Transformation (VST) using the vst function in DESeq2. The transformation was performed with blind=FALSE to account for the experimental design. The principal components of the VST transformed counts were calculated using the prcomp function in R.

### Weighted Gene Co-expression Network Analysis

WGCNA was performed using the WGCNA R package with biweight midcorrelation and a signed network topology. To reduce noise, mean absolute deviation (MAD) was calculated across all samples for each gene, and genes with MAD < 25 were excluded. Retained counts were normalized using variance-stabilizing transformation (VST) in DESeq2. Sample connectivity was assessed by computing a pairwise bicor distance adjacency matrix, followed by hierarchical clustering using the flashClust algorithm. Samples with a connectivity Z-score < −2.5 were flagged as outliers and removed prior to network construction. A soft-thresholding power was selected by using the pickSoftThreshold function of the WGCNA package. Co-expression modules were identified using blockwiseModules() with a minimum module size of 30 genes and a merge cut height of 0.25. Module colors were assigned using the labels2colors() function. Module membership scores (kME) were calculated as the Pearson correlation between each gene’s expression profile and the module eigengene, and an expanded gene set was defined for each module using a threshold of |kME| > 0.6.

### Pathway Enrichment Analysis

Pathway enrichment analysis was performed using a custom Python pipeline (v3.12.10). Modular transcriptomic datasets at each adaptive timepoint (Days 3, 7, 14, and 28) were processed using pandas (v2.1.4) and numpy (v1.26.4), from which gene symbols and corresponding log2 fold change (log2FC) values were extracted for downstream analysis. Overrepresented biological pathways were identified by querying stratified gene lists against the KEGG_2019_Mouse and Reactome_2022 reference databases using gseapy (v1.1.11), a Python interface to the Enrichr API. Gene sets with fewer than three annotated targets were excluded to minimize spurious associations. Enrichment outputs, including adjusted P values, overlapping gene identities, and overlap ratios, were aggregated across all timepoints and pivoted to align pathway terms with their respective temporal and directional conditions. The resulting matrices were ranked by minimum adjusted P value across all conditions to prioritize pathways exhibiting robust and consistent perturbation across the 28-day intervention. Pathways with an adjusted P value < 0.05 were considered statistically significant.

### UpSet Plot Visualization

Intersecting DEG sets across adaptive timepoints (Days 3, 7, 14, and 28) were computed and visualized using the upsetplot Python library. Gene identifier lists were parsed from kME-expanded modular DEG datasets using pandas, with erroneous entries removed prior to analysis. Set intersections were rendered as UpSet plots, with absolute intersection counts annotated directly on each bar. All figures were exported as vector graphics in PDF and SVG formats using matplotlib. Intersection data were additionally exported as binary membership matrices to Excel workbooks.

### Interactome Construction and Topological Cartography

To map the hierarchical architecture of the tissue-specific networks, all genes assigned to WGCNA modules were projected onto the Search Tool for the Retrieval of Interacting Genes/Proteins (STRING) database. The global interactome was constructed utilizing a confidence threshold (score ≥ 700) to ensure only empirically validated protein-protein interactions were retained. Following network assembly, topological cartography was employed to compute two spatial parameters for every node: the intra-modular connectivity (z-score), which quantifies a node’s localized density within its assigned WGCNA module, and the inter-modular participation coefficient (P), which measures the distribution of a node’s edges across all other global modules. Nodes exhibiting a z-score ≥ 2.5 were classified as topological hubs; these were further stratified by their pleiotropic distribution into provincial hubs (P < 0.30), connector hubs (0.30 ≤ P < 0.75), and kinless hubs (P≥0.75).

### Extraction of Systemic Master Initiators

To identify the primary regulatory drivers orchestrating early tissue remodeling, the topological matrix was filtered to isolate hubs exhibiting significant differential expression exclusively during the acute transition phases (Day 3 and Day 7). The analysis was strictly constrained to connector and kinless hubs, as their elevated participation coefficients denote extensive geometric outreach across distinct co-expression modules. The resulting candidates were designated as early systemic initiators and extracted for localized neighborhood mapping.

### Topological Network Computation

To model the global and isolated regulatory topologies, spatial coordinate matrices were derived utilizing the Fruchterman-Reingold force-directed algorithm. Master network architectures were subsequently computed to track the discrete temporal activation states of the initiator cascade across the 28-day paradigm, isolating the active master initiators and their first-degree downstream targets from the quiescent background interactome.

### Functional Enrichment Profiling

To delineate the specific biochemical and signaling pathways orchestrated by the early initiator cascade, the master hubs and their immediate first-degree interactome neighbors were subjected to targeted functional enrichment utilizing the GSEAPY interface. Hypergeometric statistical testing was executed in parallel across the Gene Ontology (Biological Process), KEGG, and Reactome databases. Pathways satisfying a strict false discovery rate threshold (FDR < 0.05) were retained and hierarchically sorted by their maximum -log_10_(P_adj_) values to systematically extract the dominant metabolic and signaling cascades governing the temporal response.

### Hierarchically Clustered Dual-Tissue Heatmap

Differentially expressed transcripts derived from liver and inguinal white adipose tissue (days 3, 7, 14, and 28; adjusted *P* < 0.05, |LFC| ≥ 1.0). were assembled into a matrix and subjected to unsupervised hierarchical clustering (Euclidean distance, Ward’s linkage); the resulting dendrogram was partitioned into eight modules. To evaluate temporal expression dynamics, a consensus eigengene was computed for each module by calculating the mean log-fold change across its constituent members at each discrete time point.

### Intersection with Longevity-Associated Gene Databases

To assess the overlap between these empirically derived co-expression modules and validated longevity-associated loci, the identified transcripts were cross-referenced against the GenAge database (Human Ageing Genomic Resources). This comparison encompassed both human cohorts and established model organisms (Saccharomyces cerevisiae, Caenorhabditis elegans, Drosophila melanogaster, and Mus musculus). Intersecting transcripts were subsequently annotated with Boolean identifiers specifying their association with human aging or model organism lifespan.

### Principal Component Analysis biplot

To quantify the structural reconfiguration of these overarching regulatory networks, multivariate dimensionality reduction was executed via Principal Component Analysis (scikit-learn). The phenotypic matrix was standardized utilizing a z-score transformation, ensuring that physiological parameters with disparate scalar units contributed equivalently to the covariance matrix. A global eigenspace was defined utilizing the comprehensive 28-day dataset, establishing fixed orthogonal axes driven by the primary components of longitudinal variance (PC1 and PC2). Following the extraction of normalized phenotypic loadings, the independent temporal transcriptomic matrices were projected directly onto this master coordinate system. To determine the structural dominance of individual transcripts within the regulatory hierarchy, Pearson correlation coefficients (r) were calculated against the global sample scores; the magnitude of each transcript was defined by its Euclidean vector length from the geometric origin 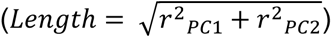. The transcriptomic matrix was then filtered to isolate the upper 5% of targets exhibiting the highest cumulative Euclidean magnitudes, delineating the primary adaptive network. Lastly, aggregate structural magnitudes were computed across the temporal span to isolate the 20 universal master regulators; their functional polarities were delineated by the dot product between transcript coordinates and the composite phenotypic vector, classifying targets as either metabolic activators or suppressors of the intervention-induced state.

### Data Processing and Statistical Analysis

Data processing and statistical analyses were performed using custom scripts written in Python (v3.12.10) unless otherwise specified. Group comparisons for metabolic and physiological data were performed using two-way analysis of variance (ANOVA) multiple comparisons test with Šidák’s post hoc test. Data are presented as mean ± s.e.m. unless otherwise stated. Statistical significance was defined as P < 0.05. Sample sizes are reported in the figure legends. Energy expenditure data were analyzed using Analysis of Covariance (ANCOVA) to adjust for differences in body mass. Body weight served as the continuous covariate, and feeding group (AL vs. CR) served as the independent fixed factor. Prior to testing group differences, the homogeneity of regression slopes was assessed by modeling the interaction term between Group and Body Weight (Group × Weight). In the absence of significant interaction (*p* > 0.05), a standard ANCOVA model was applied to test for the main effect of Group. Statistical calculations were performed using the statsmodels and pingouin libraries.

## Acknowledgements

The laboratory of J.S.T. is supported in part by the National Institutes of Health (NIH)/NIA R01AG072736, NIH/NIDDK R01 DK140283, NIH/NINDS R01 NS114527, and The Milky Way Research Foundation. H.H.P. is supported in part by Life Science Research Foundation sponsored by the Welch Foundation R-I-0001-20230405 and American Federation for Aging Research-Glenn Foundation Postdoctoral Fellowship. J.S.T. was supported as an Investigator in the Howard Hughes Medical Institute.

---

**Extended Data Figure 1.**
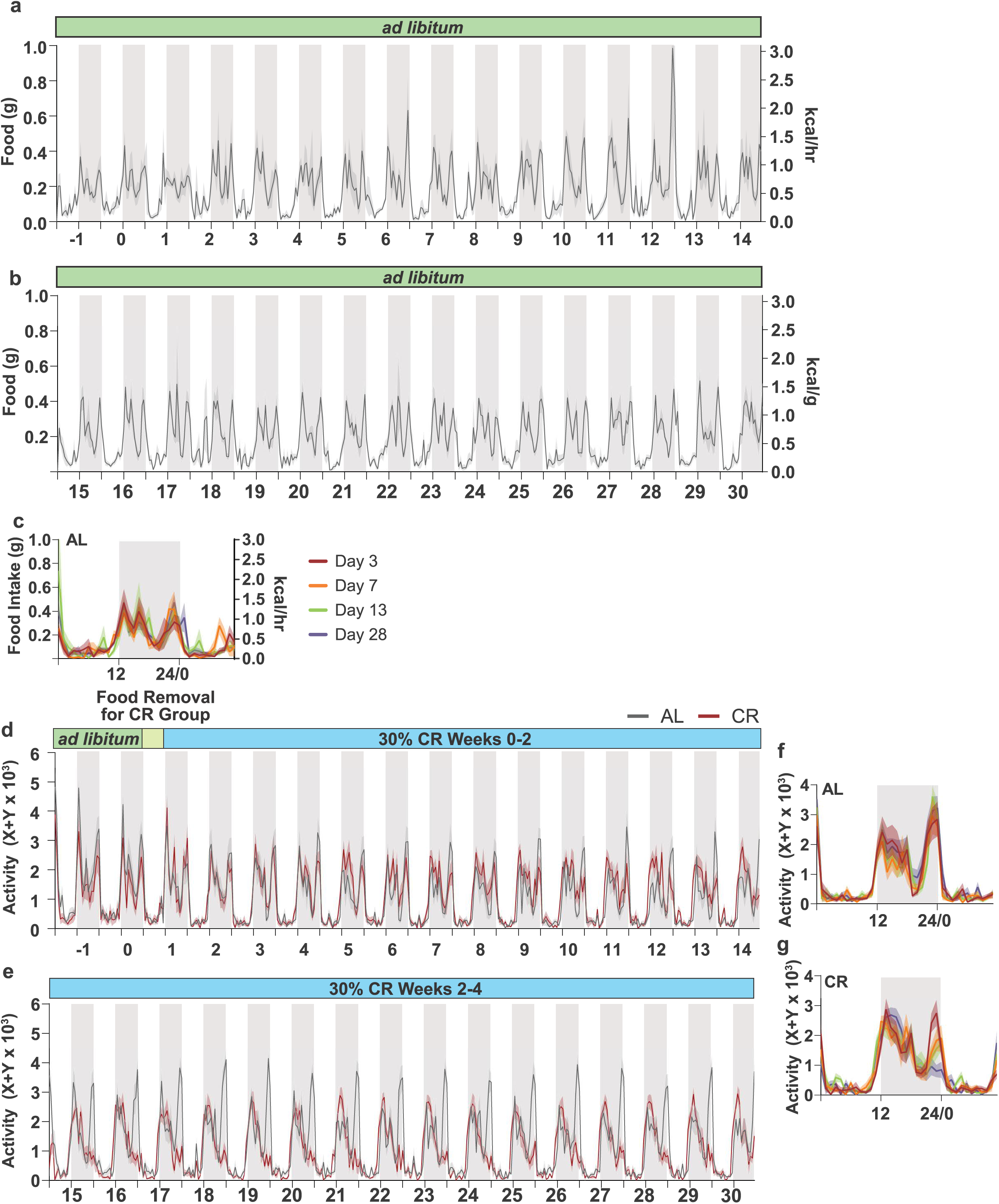
Locomotor activity and food consumption profiles across the four-week metabolic chamber recording period. **a, b,** Daily food consumption versus time for AL groups only. **c,** Overlay of food intake activity for AL at Days 3, 7, 13, and 28. **d, e,** Ambulatory activity versus time for AL and CR cohorts. **f, g,** Overlay of locomoter activity for AL and CR at Days 3, 7, 13, and 28. Shaded regions denote the dark cycle. Data are mean± s.e.m.; n = 10-12 per group. **Related to** Figures 1-2.

**Extended Data Figure 2.**
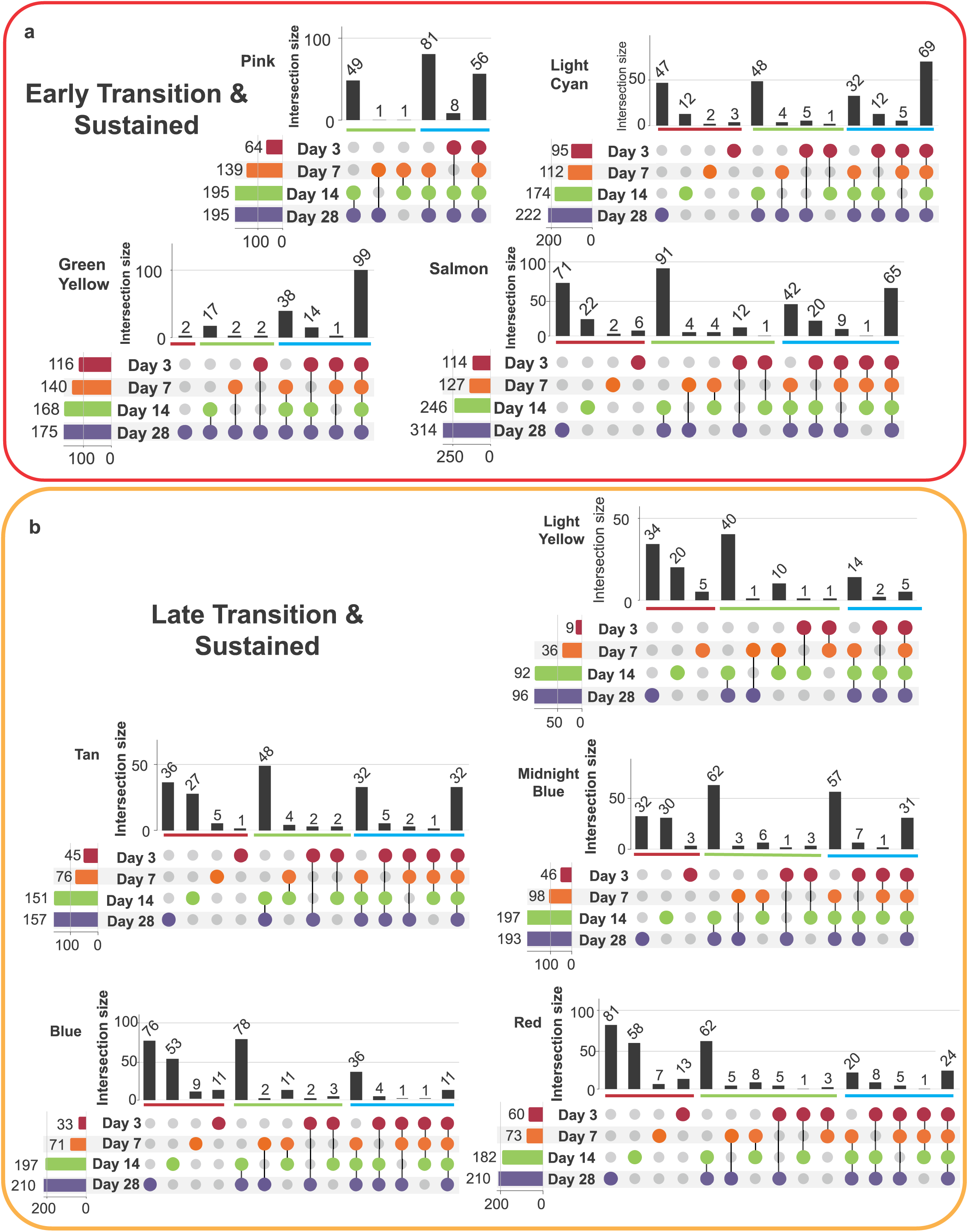
UpSet analysis of hepatic DEGs resolves early transition-sustained and late transition-sustained clusters. UpSet plots for the **a,** Early Transition-Sustained cluster (Green Yellow, Pink, Light Cyan, and Salmon modules) and **b,** Late Transition-Sustained cluster (Tan, Blue, Midnight Blue, Red, and Light Yellow modules). DEGs defined as adjusted P < 0.05 from kME-expanded gene sets (lkMEI > 0.6). Horizontal bars indicate total DEG count per adaptive timepoint; vertical bars indicate the size of intersecting gene sets. **Related to** Figure 4.

**Extended Data Figure 3.**
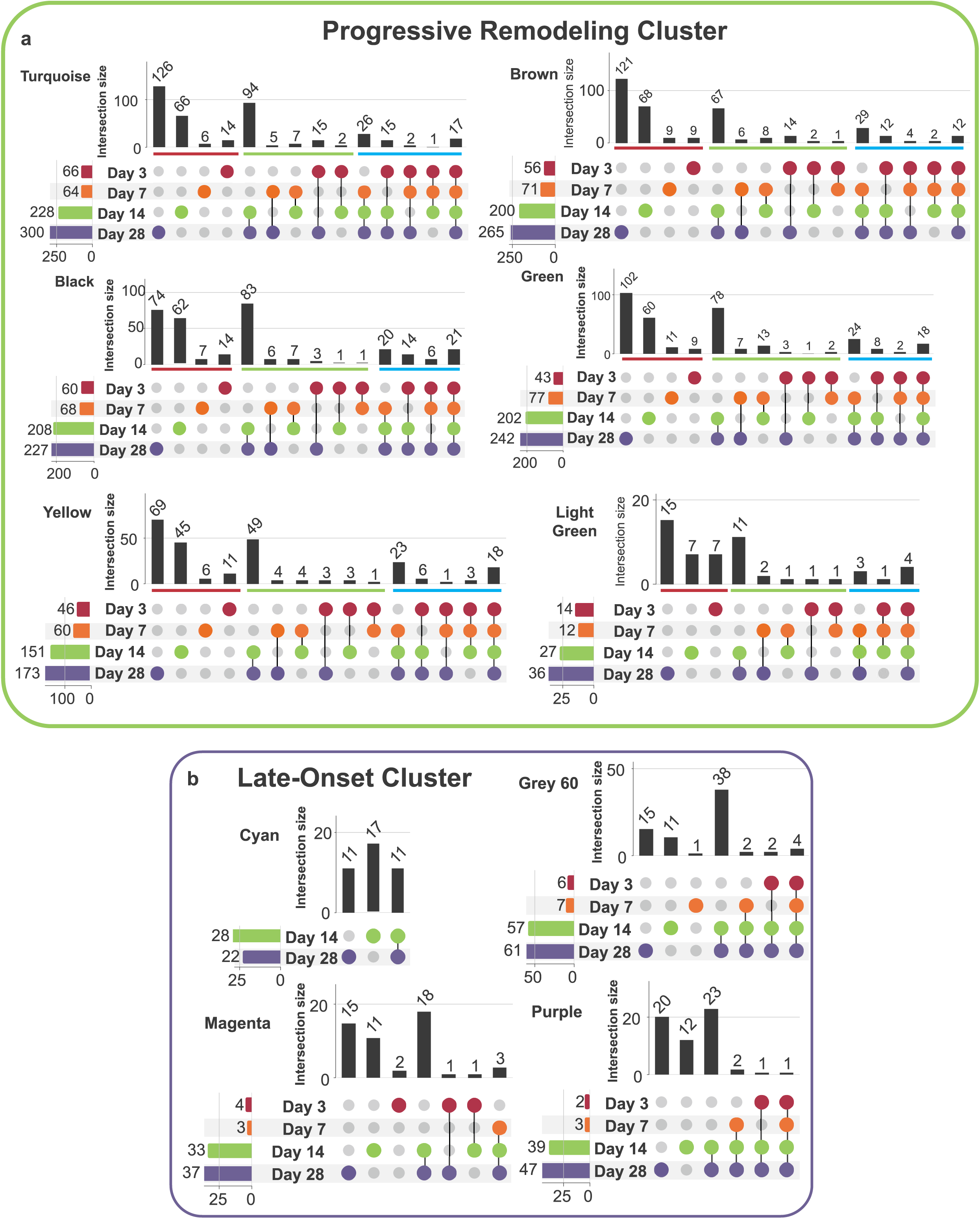
UpSet analysis of hepatic DEGs resolves progressive remodeling and late-onset clusters. UpSet plots for the **a,** Progressive Remodeling cluster (Turquoise, Brown, Black, Green, Yellow, and Light Green modules) and **b,** Late-Onset cluster (Grey60, Cyan, Magenta, and Purple modules). DEGs defined as adjusted P < 0.05 from kME-expanded gene sets (lkMEI > 0.6). Horizontal bars indicate total DEG count per adaptive timepoint; vertical bars indicate the size of intersecting gene sets. **Related to** Figure 4.

**Extended Data Fig. 4.**
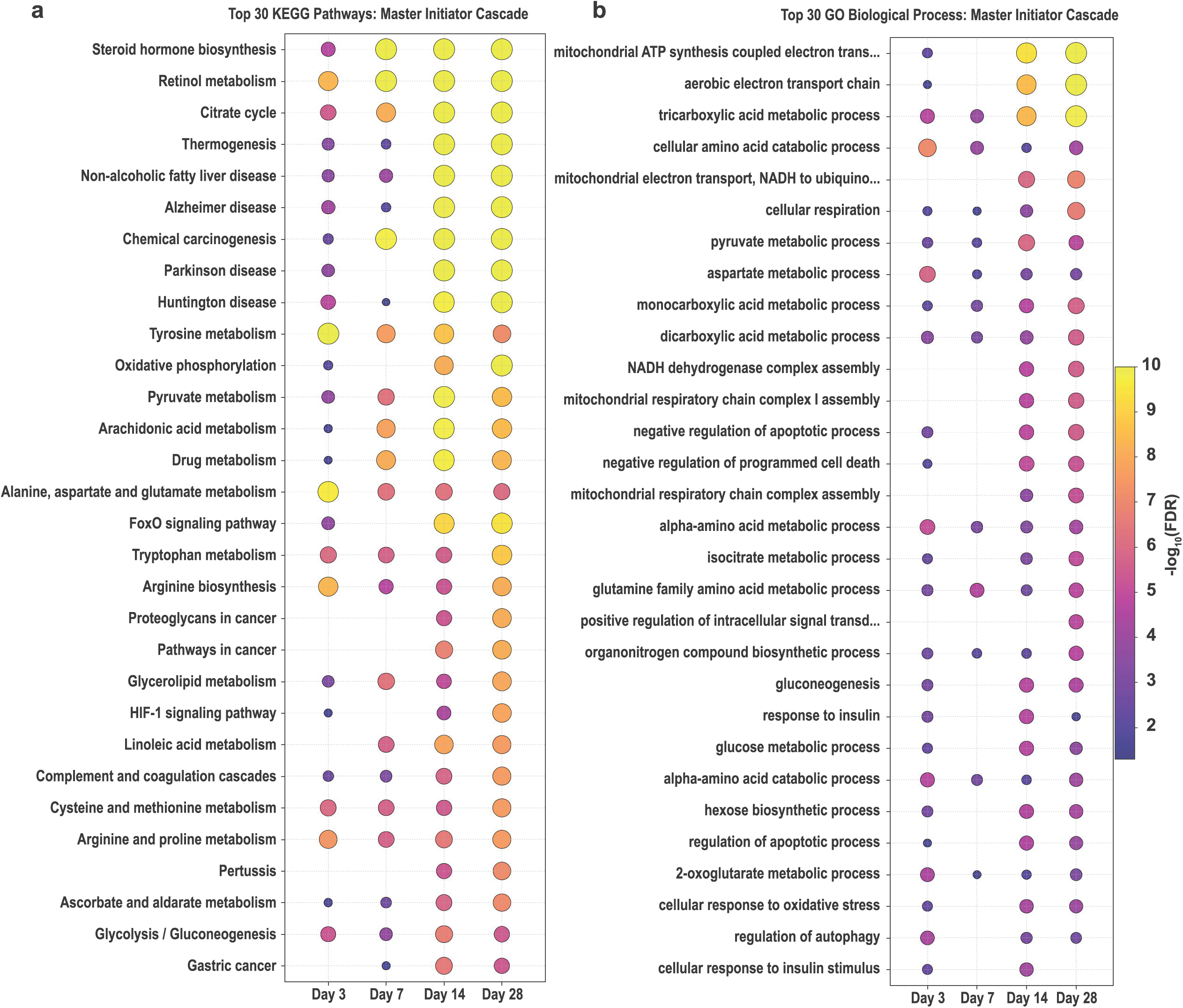
Top 30 KEGG and GO enrichment analyses of the hepatic initiator network. **a,** KEGG pathway enrichment of the consolidated hepatic hub network. **b,** GO Biological Process enrichment. **Related to** Figure 5.

**Extended Data Figure 5.**
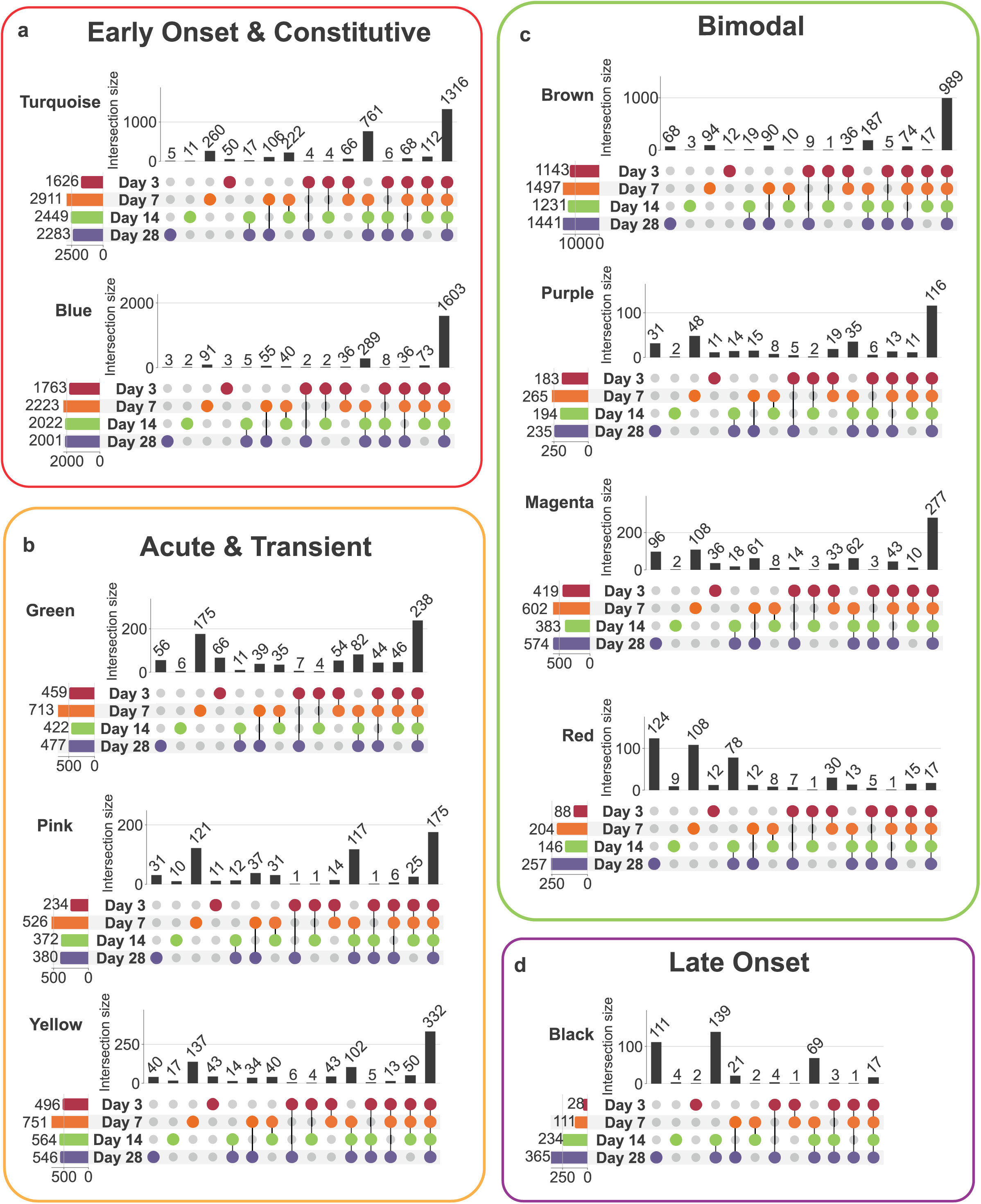
UpSet analysis of iWAT DEGs resolves four adaptation clusters. UpSet plots for the **a,** Early-Onset Constitutive cluster (Blue and Turquoise modules). **b,** Acute Transient cluster (Green, Pink, and Yellow modules). **c,** Non-Monotonic cluster (Brown, Purple, Magenta, and Red modules). **d,** Late-Onset cluster (Black module). DEGs defined as adjusted P < 0.05 from kME-expanded gene sets (lkMEI > 0.6). Horizontal bars indicate total DEG count per adaptive timepoint; vertical bars indicate the size of intersecting gene sets. **Related to** Figure 6.

**Extended Data Fig. 6.**
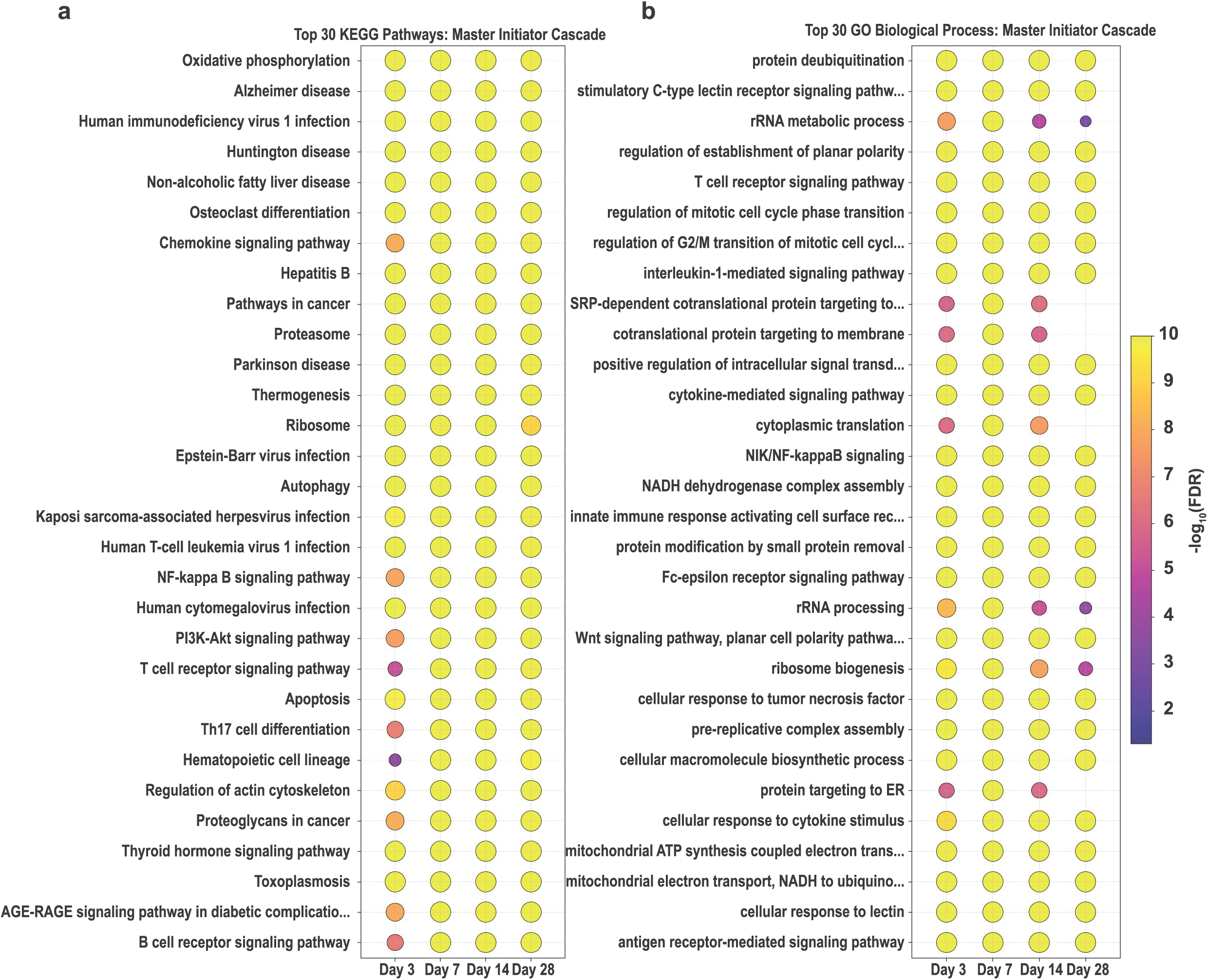
Top 30 KEGG and GO enrichment analyses of the iWAT initiator network. **a,** KEGG pathway enrichment of the consolidated hepatic hub network. **b,** GO Biological Process enrichment. **Related to** Figure 7.

**Extended Data Figs. 7.**
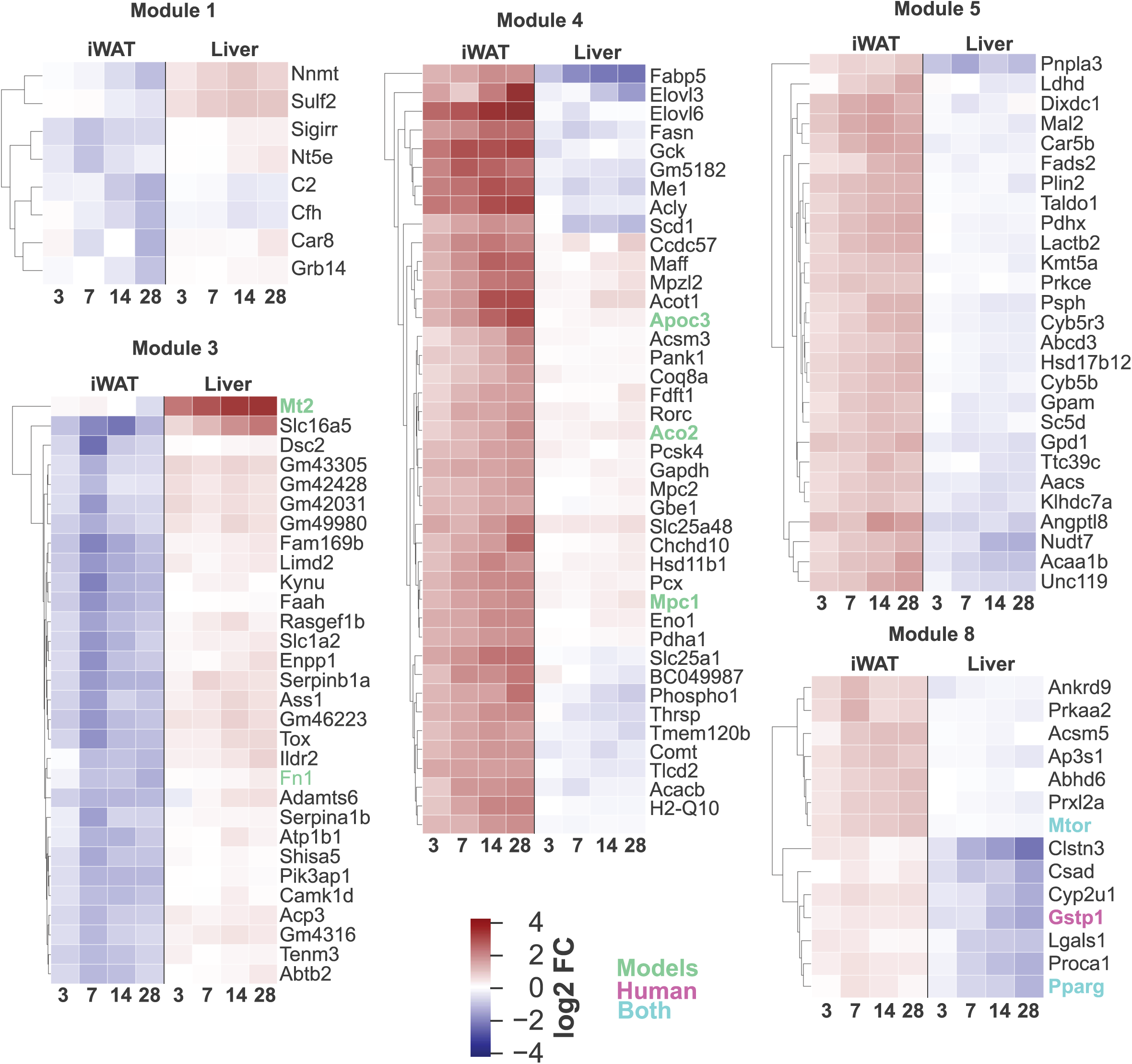
Cross-tissue hierarchical clustering of shared CR-regulated transcripts. Heatmaps of the 1,232 transcripts regulated in both liver and iWAT across the adaptive timeline, filtered to ILFCI 2::: 1.0 and organized by unsupervised hierarchical clustering into eight modules. Modules with divergent tissue-specific expression trajectories: 1, 3, 4, 5, and 8. Module eigengene profiles are shown in **Supplementary Fig. 8**.

**Extended Data Figs. 8.**
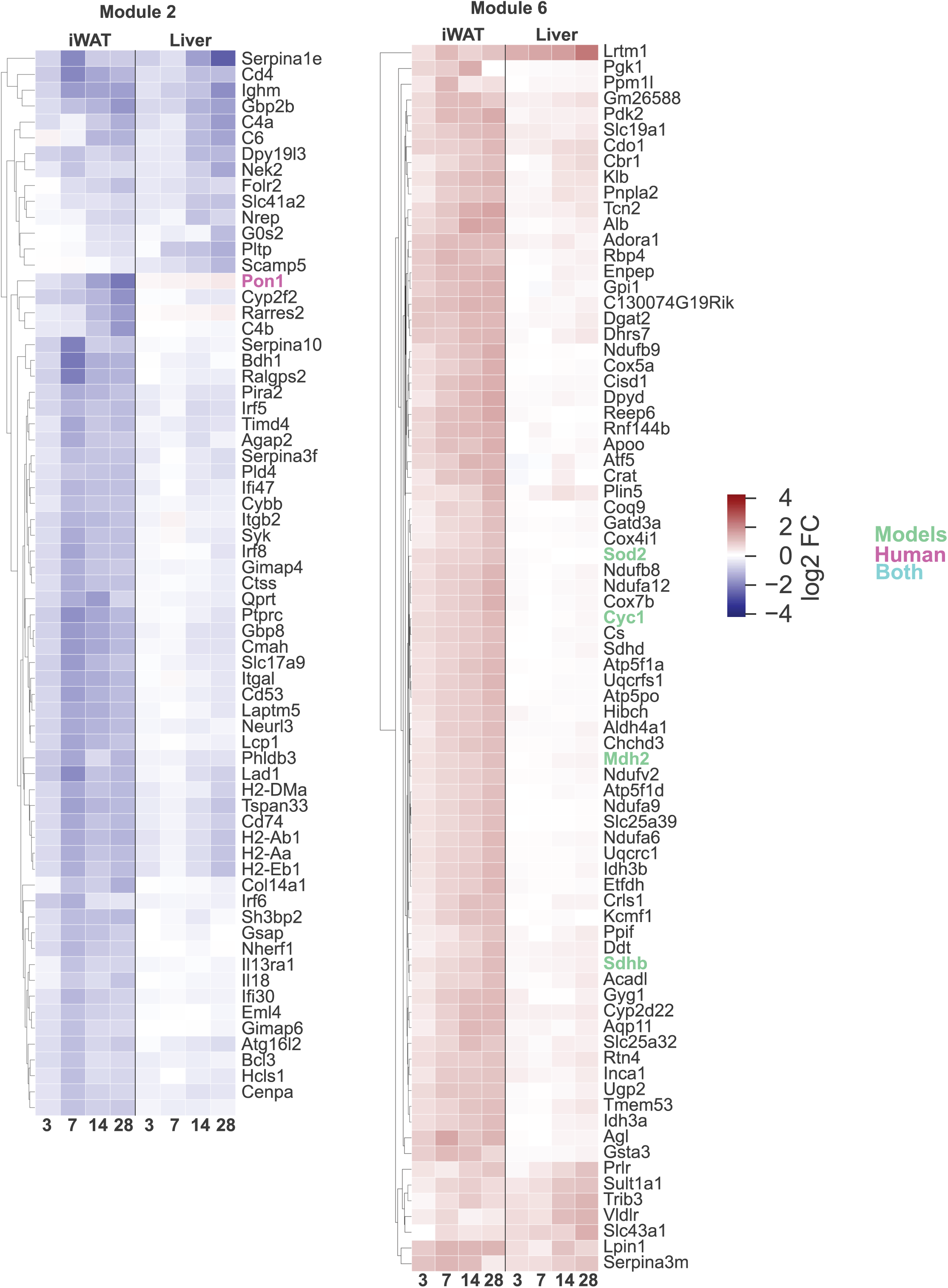
Cross-tissue hierarchical clustering of shared CR-regulated transcripts. Heatmaps of the 1,232 transcripts regulated in both liver and iWAT across the adaptive timeline, filtered to ILFCI 2::: 1.0 and organized by unsupervised hierarchical clustering into eight modules. Modules with concordant tissue-specific expression trajectories: 2 and 6. Longevity-associated genes from the GenAge database are highlighted. Module eigengene profiles are shown in Supplementary Fig. 8.

## Notes

### Competing Interest Statement

The authors have declared no competing interest.

## References

1 Colman, R. J. et al. Caloric restriction delays disease onset and mortality in rhesus monkeys. Science 325, 201–204 (2009). 10.1126/science.1173635

2 Lin, S. J., Defossez, P. A. & Guarente, L. Requirement of NAD and SIR2 for life-span extension by calorie restriction in Saccharomyces cerevisiae. Science 289, 2126–2128 (2000). 10.1126/science.289.5487.2126

3 Weindruch, R., Walford, R. L., Fligiel, S. & Guthrie, D. The retardation of aging in mice by dietary restriction: longevity, cancer, immunity and lifetime energy intake. J Nutr 116, 641–654 (1986). 10.1093/jn/116.4.641

4 Lakowski, B. & Hekimi, S. The genetics of caloric restriction in Caenorhabditis elegans. Proceedings of the National Academy of Sciences 95, 13091–13096 (1998). 10.1073/pnas.95.22.13091

5 McCay, C. M., Crowell, M. F. & Maynard, L. A. The Effect of Retarded Growth Upon the Length of Life Span and Upon the Ultimate Body Size: One Figure. The Journal of Nutrition 10, 63–79 (1935). 10.1093/jn/10.1.63

6 Taormina, G. & Mirisola, M. G. Calorie restriction in mammals and simple model organisms. Biomed Res Int 2014, 308690 (2014). 10.1155/2014/308690

7 Meydani, S. N. et al. Long-term moderate calorie restriction inhibits inflammation without impairing cell-mediated immunity: a randomized controlled trial in non-obese humans. Aging (Albany NY) 8, 1416–1431 (2016). 10.18632/aging.100994

8 Lopez-Lluch, G. et al. Calorie restriction induces mitochondrial biogenesis and bioenergetic efficiency. Proc Natl Acad Sci U S A 103, 1768–1773 (2006). 10.1073/pnas.0510452103

9 Fontana, L., Meyer, T. E., Klein, S. & Holloszy, J. O. Long-term calorie restriction is highly effective in reducing the risk for atherosclerosis in humans. Proc Natl Acad Sci U S A 101, 6659–6663 (2004). 10.1073/pnas.0308291101

10 Colman, R. J. et al. Caloric restriction reduces age-related and all-cause mortality in rhesus monkeys. Nature communications 5, 3557 (2014). 10.1038/ncomms4557

11 Civitarese, A. E. et al. Calorie restriction increases muscle mitochondrial biogenesis in healthy humans. PLoS Med 4, e76 (2007). 10.1371/journal.pmed.0040076

12 Barzilai, N., Banerjee, S., Hawkins, M., Chen, W. & Rossetti, L. Caloric restriction reverses hepatic insulin resistance in aging rats by decreasing visceral fat. J Clin Invest 101, 1353–1361 (1998). 10.1172/JCI485

13 Selman, C. et al. Ribosomal protein S6 kinase 1 signaling regulates mammalian life span. Science 326, 140–144 (2009). 10.1126/science.1177221

14 Holzenberger, M. et al. IGF-1 receptor regulates lifespan and resistance to oxidative stress in mice. Nature 421, 182–187 (2003). 10.1038/nature01298

15 Canto, C. et al. AMPK regulates energy expenditure by modulating NAD+ metabolism and SIRT1 activity. Nature 458, 1056–1060 (2009). 10.1038/nature07813

16 Bonkowski, M. S., Rocha, J. S., Masternak, M. M., Al Regaiey, K. A. & Bartke, A. Targeted disruption of growth hormone receptor interferes with the beneficial actions of calorie restriction. Proc Natl Acad Sci U S A 103, 7901–7905 (2006). 10.1073/pnas.0600161103

17 Boily, G. et al. SirT1 regulates energy metabolism and response to caloric restriction in mice. PLoS One 3, e1759 (2008). 10.1371/journal.pone.0001759

18 Bjedov, I. et al. Mechanisms of life span extension by rapamycin in the fruit fly Drosophila melanogaster. Cell metabolism 11, 35–46 (2010). 10.1016/j.cmet.2009.11.010

19 Apfeld, J., O’Connor, G., McDonagh, T., DiStefano, P. S. & Curtis, R. The AMP-activated protein kinase AAK-2 links energy levels and insulin-like signals to lifespan in C. elegans. Genes Dev 18, 3004–3009 (2004). 10.1101/gad.1255404

20 Cohen, H. Y. et al. Calorie restriction promotes mammalian cell survival by inducing the SIRT1 deacetylase. Science 305, 390–392 (2004). 10.1126/science.1099196

21 Um, S. H. et al. Absence of S6K1 protects against age- and diet-induced obesity while enhancing insulin sensitivity. Nature 431, 200–205 (2004). 10.1038/nature02866

22 Tremblay, F. et al. Identification of IRS-1 Ser-1101 as a target of S6K1 in nutrient- and obesity-induced insulin resistance. Proc Natl Acad Sci U S A 104, 14056–14061 (2007). 10.1073/pnas.0706517104

23 O’Reilly, K. E. et al. mTOR inhibition induces upstream receptor tyrosine kinase signaling and activates Akt. Cancer Res 66, 1500–1508 (2006). 10.1158/0008-5472.CAN-05-2925

24 Mouchiroud, L. et al. The NAD(+)/Sirtuin Pathway Modulates Longevity through Activation of Mitochondrial UPR and FOXO Signaling. Cell 154, 430–441 (2013). 10.1016/j.cell.2013.06.016

25 Jaafar, R. et al. mTORC1 to AMPK switching underlies beta-cell metabolic plasticity during maturation and diabetes. J Clin Invest 129, 4124–4137 (2019). 10.1172/JCI127021

26 Harrington, L. S. et al. The TSC1-2 tumor suppressor controls insulin-PI3K signaling via regulation of IRS proteins. J Cell Biol 166, 213–223 (2004). 10.1083/jcb.200403069

27 Carracedo, A. et al. Inhibition of mTORC1 leads to MAPK pathway activation through a PI3K-dependent feedback loop in human cancer. J Clin Invest 118, 3065–3074 (2008). 10.1172/JCI34739

28 Lan, F., Cacicedo, J. M., Ruderman, N. & Ido, Y. SIRT1 modulation of the acetylation status, cytosolic localization, and activity of LKB1. Possible role in AMP-activated protein kinase activation. J Biol Chem 283, 27628–27635 (2008). 10.1074/jbc.M805711200

29 Sato, S. et al. Circadian Reprogramming in the Liver Identifies Metabolic Pathways of Aging. Cell 170, 664-677 e611 (2017). 10.1016/j.cell.2017.07.042

30 Acosta-Rodriguez, V. et al. Circadian alignment of early onset caloric restriction promotes longevity in male C57BL/6J mice. Science 376, 1192–1202 (2022). 10.1126/science.abk0297

31 Pletcher, S. D. et al. Genome-wide transcript profiles in aging and calorically restricted Drosophila melanogaster. Curr Biol 12, 712–723 (2002). 10.1016/s0960-9822(02)00808-4

32 Lee, C. K., Klopp, R. G., Weindruch, R. & Prolla, T. A. Gene expression profile of aging and its retardation by caloric restriction. Science 285, 1390–1393 (1999). 10.1126/science.285.5432.1390

33 Kayo, T., Allison, D. B., Weindruch, R. & Prolla, T. A. Influences of aging and caloric restriction on the transcriptional profile of skeletal muscle from rhesus monkeys. Proc Natl Acad Sci U S A 98, 5093–5098 (2001). 10.1073/pnas.081061898

34 Cao, S. X., Dhahbi, J. M., Mote, P. L. & Spindler, S. R. Genomic profiling of short- and long-term caloric restriction effects in the liver of aging mice. Proc Natl Acad Sci U S A 98, 10630–10635 (2001). 10.1073/pnas.191313598

35 Acosta-Rodriguez, V. A., de Groot, M. H. M., Rijo-Ferreira, F., Green, C. B. & Takahashi, J. S. Mice under Caloric Restriction Self-Impose a Temporal Restriction of Food Intake as Revealed by an Automated Feeder System. Cell metabolism 26, 267–277 e262 (2017). 10.1016/j.cmet.2017.06.007

36 Pak, H. H. et al. Non-canonical metabolic and molecular effects of calorie restriction are revealed by varying temporal conditions. Cell reports 43, 114663 (2024). 10.1016/j.celrep.2024.114663

37 Pak, H. H. et al. Fasting drives the metabolic, molecular and geroprotective effects of a calorie-restricted diet in mice. Nat Metab 3, 1327–1341 (2021). 10.1038/s42255-021-00466-9

38 Bruss, M. D., Khambatta, C. F., Ruby, M. A., Aggarwal, I. & Hellerstein, M. K. Calorie restriction increases fatty acid synthesis and whole body fat oxidation rates. Am J Physiol Endocrinol Metab 298, E108–116 (2010). 10.1152/ajpendo.00524.2009

39 Guimera, R. & Nunes Amaral, L. A. Functional cartography of complex metabolic networks. Nature 433, 895–900 (2005). 10.1038/nature03288

40 Rosen, E. D. & Spiegelman, B. M. Adipocytes as regulators of energy balance and glucose homeostasis. Nature 444, 847–853 (2006). 10.1038/nature05483

41 Lee, D. E., Kehlenbrink, S., Lee, H., Hawkins, M. & Yudkin, J. S. Getting the message across: mechanisms of physiological cross talk by adipose tissue. Am J Physiol Endocrinol Metab 296, E1210–1229 (2009). 10.1152/ajpendo.00015.2009

42 Kim, J. B. Dynamic cross talk between metabolic organs in obesity and metabolic diseases. Exp Mol Med 48, e214 (2016). 10.1038/emm.2015.119

43 Azzu, V., Vacca, M., Virtue, S., Allison, M. & Vidal-Puig, A. Adipose Tissue-Liver Cross Talk in the Control of Whole-Body Metabolism: Implications in Nonalcoholic Fatty Liver Disease. Gastroenterology 158, 1899–1912 (2020). 10.1053/j.gastro.2019.12.054

44 Luo, L. & Liu, M. Adipose tissue in control of metabolism. J Endocrinol 231, R77–R99 (2016). 10.1530/JOE-16-0211

45 Choe, S. S., Huh, J. Y., Hwang, I. J., Kim, J. I. & Kim, J. B. Adipose Tissue Remodeling: Its Role in Energy Metabolism and Metabolic Disorders. Front Endocrinol (Lausanne) 7, 30 (2016). 10.3389/fendo.2016.00030

46 Yang, A. & Mottillo, E. P. Adipocyte lipolysis: from molecular mechanisms of regulation to disease and therapeutics. Biochem J 477, 985–1008 (2020). 10.1042/BCJ20190468

47 Zhang, S. et al. Short-term moderate caloric restriction in a high-fat diet alleviates obesity via AMPK/SIRT1 signaling in white adipocytes and liver. Food & nutrition research 66 (2022). 10.29219/fnr.v66.7909

48 Shaw, R. J. LKB1 and AMP-activated protein kinase control of mTOR signalling and growth. Acta Physiol (Oxf) 196, 65–80 (2009). 10.1111/j.1748-1716.2009.01972.x

49 Canto, C. & Auwerx, J. Calorie restriction: is AMPK a key sensor and effector? Physiology (Bethesda) 26, 214–224 (2011). 10.1152/physiol.00010.2011

50 Giacomello, E. & Toniolo, L. The Potential of Calorie Restriction and Calorie Restriction Mimetics in Delaying Aging: Focus on Experimental Models. Nutrients 13 (2021). 10.3390/nu13072346

51 Tulsian, R., Velingkaar, N. & Kondratov, R. Caloric restriction effects on liver mTOR signaling are time-of-day dependent. Aging (Albany NY) 10, 1640–1648 (2018). 10.18632/aging.101498

52 Mitchell, S. E. et al. The effects of graded levels of calorie restriction: I. impact of short term calorie and protein restriction on body composition in the C57BL/6 mouse. Oncotarget 6, 15902–15930 (2015). 10.18632/oncotarget.4142

53 de Cabo, R. & Mattson, M. P. Effects of Intermittent Fasting on Health, Aging, and Disease. N Engl J Med 381, 2541–2551 (2019). 10.1056/NEJMra1905136

54 Chaix, A., Lin, T., Le, H. D., Chang, M. W. & Panda, S. Time-Restricted Feeding Prevents Obesity and Metabolic Syndrome in Mice Lacking a Circadian Clock. Cell metabolism 29, 303–319 e304 (2019). 10.1016/j.cmet.2018.08.004

55 Mitchell, S. J. et al. Daily Fasting Improves Health and Survival in Male Mice Independent of Diet Composition and Calories. Cell metabolism 29, 221–228 e223 (2019). 10.1016/j.cmet.2018.08.011

56 Deota, S. et al. Diurnal transcriptome landscape of a multi-tissue response to time-restricted feeding in mammals. Cell metabolism 35, 150–165 e154 (2023). 10.1016/j.cmet.2022.12.006

57 Defour, M., Hooiveld, G., van Weeghel, M. & Kersten, S. Probing metabolic memory in the hepatic response to fasting. Physiol Genomics 52, 602–617 (2020). 10.1152/physiolgenomics.00117.2020

58 Goldstein, I. et al. Transcription factor assisted loading and enhancer dynamics dictate the hepatic fasting response. Genome Res 27, 427–439 (2017). 10.1101/gr.212175.116

59 Sokolovic, M. et al. The transcriptomic signature of fasting murine liver. BMC Genomics 9, 528 (2008). 10.1186/1471-2164-9-528

60 Li, F. et al. Apoptotic cells activate the “phoenix rising” pathway to promote wound healing and tissue regeneration. Science signaling 3, ra13 (2010). 10.1126/scisignal.2000634

61 Wilson, C. H. & Kumar, S. Caspases in metabolic disease and their therapeutic potential. Cell death and differentiation 25, 1010–1024 (2018). 10.1038/s41418-018-0111-x

62 Eskandari, E. & Eaves, C. J. Paradoxical roles of caspase-3 in regulating cell survival, proliferation, and tumorigenesis. J Cell Biol 221 (2022). 10.1083/jcb.202201159

63 Cao, Z. et al. Sublethal executioner caspase activation in hepatocytes promotes liver regeneration through the JAK/STAT3 pathway. PLoS Biol 23, e3003357 (2025). 10.1371/journal.pbio.3003357

64 Barthelemy, J., Bogard, G. & Wolowczuk, I. Beyond energy balance regulation: The underestimated role of adipose tissues in host defense against pathogens. Front Immunol 14, 1083191 (2023). 10.3389/fimmu.2023.1083191

65 Brestoff, J. R. et al. Group 2 innate lymphoid cells promote beiging of white adipose tissue and limit obesity. Nature 519, 242–246 (2015). 10.1038/nature14115

66 Fabbiano, S. et al. Caloric Restriction Leads to Browning of White Adipose Tissue through Type 2 Immune Signaling. Cell metabolism 24, 434–446 (2016). 10.1016/j.cmet.2016.07.023

67 Fernandes, M. et al. Systematic analysis of the gerontome reveals links between aging and age-related diseases. Hum Mol Genet 25, 4804–4818 (2016). 10.1093/hmg/ddw307

68 Swindell, W. R. Genes and gene expression modules associated with caloric restriction and aging in the laboratory mouse. BMC Genomics 10, 585 (2009). 10.1186/1471-2164-10-585

69 Barger, J. L. et al. Identification of tissue-specific transcriptional markers of caloric restriction in the mouse and their use to evaluate caloric restriction mimetics. Aging cell 16, 750–760 (2017). 10.1111/acel.12608

70 Miller, R. A. et al. Adiponectin suppresses gluconeogenic gene expression in mouse hepatocytes independent of LKB1-AMPK signaling. J Clin Invest 121, 2518–2528 (2011). 10.1172/JCI45942

71 Ruan, H. & Dong, L. Q. Adiponectin signaling and function in insulin target tissues. J Mol Cell Biol 8, 101–109 (2016). 10.1093/jmcb/mjw014

72 Li, X. et al. Mechanisms by which adiponectin reverses high fat diet-induced insulin resistance in mice. Proc Natl Acad Sci U S A 117, 32584–32593 (2020). 10.1073/pnas.1922169117

73 Yamauchi, T. et al. Adiponectin stimulates glucose utilization and fatty-acid oxidation by activating AMP-activated protein kinase. Nat Med 8, 1288–1295 (2002). 10.1038/nm788

