## Supplementary Figure for "Two Stages of Dynamic Metabolic and Transcriptomic Remodeling During the Adaptation to Caloric Restriction in Male C57BL/6J"

### Supplementary Figure 1

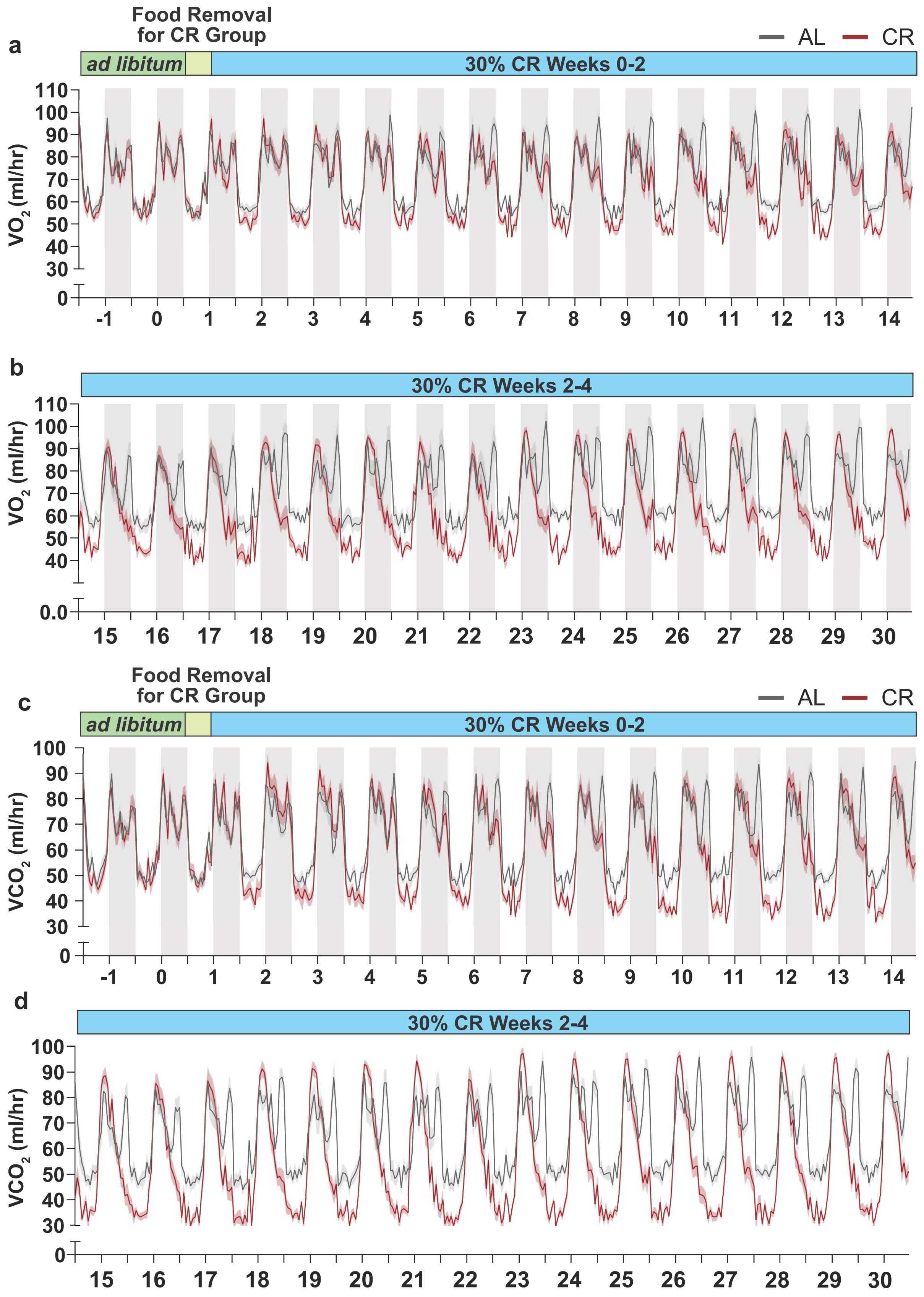

**Supplementary Figure 1. Whole-body $VO_2$ and $VCO_2$ profiles across the four-week recording period.** **a**, $VO_2$ profiles for AL and CR cohorts stratified by adaptive timepoint. **b**, $VCO_2$ profiles for AL and CR cohorts stratified by adaptive timepoint. Shaded regions denote the dark cycle. Data are mean ± s.e.m.; n = 10-12 biologically independent animals per group. Related to Figure 1.

### Supplementary Figure 2

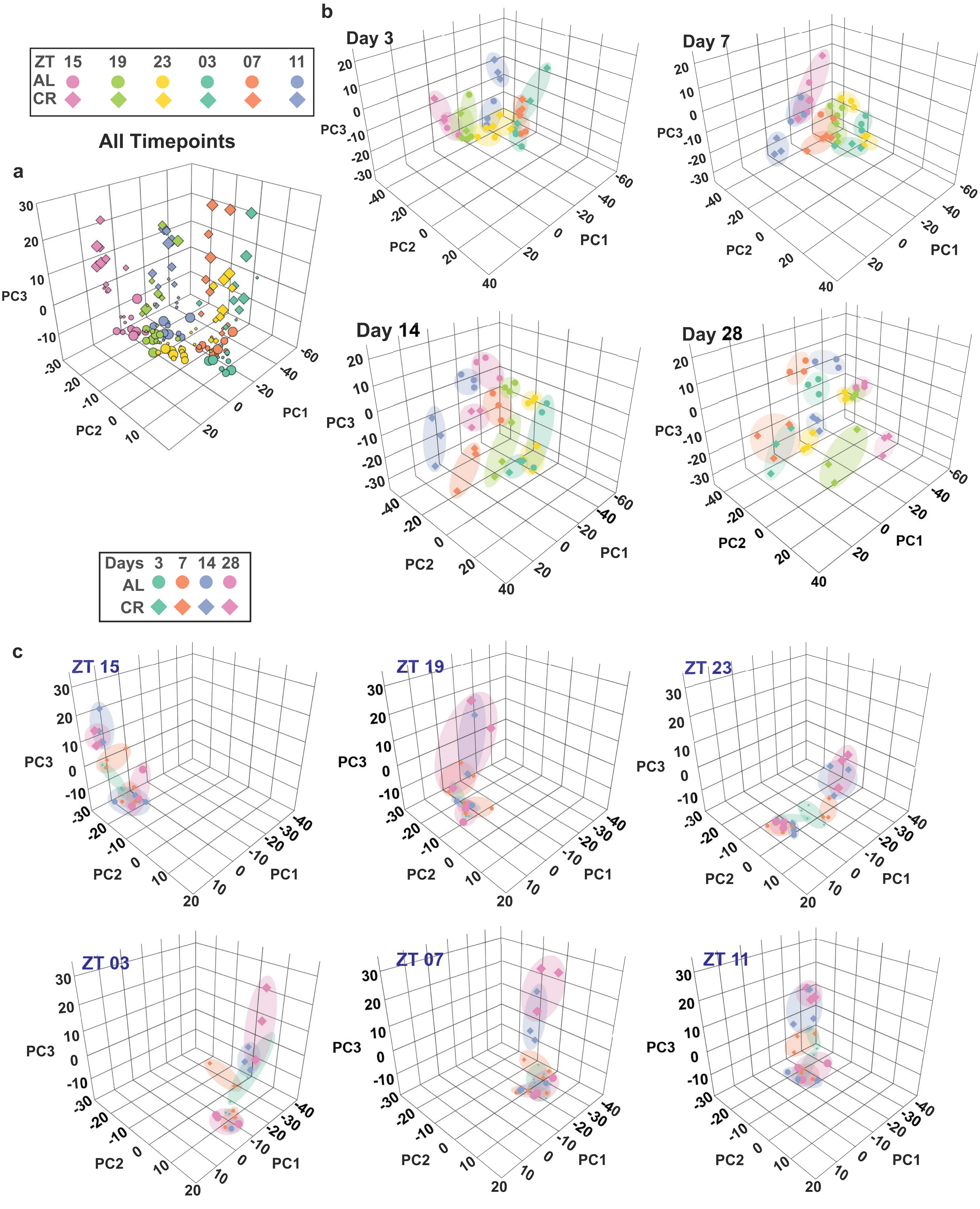

**Supplementary Figure 2. Principal component analysis of the hepatic transcriptome.** **a**, PCA stratified by dietary condition (AL vs. CR). **b**, PCA stratified by adaptive timepoint. **c**, PCA stratified by time of day. Data represent normalized transcript counts from bulk RNA sequencing. n = 3 biological independent mice per feeding condition per adaptive timepoint per ZT timepoints. Related to Figure 4.

### Supplementary Figure 3

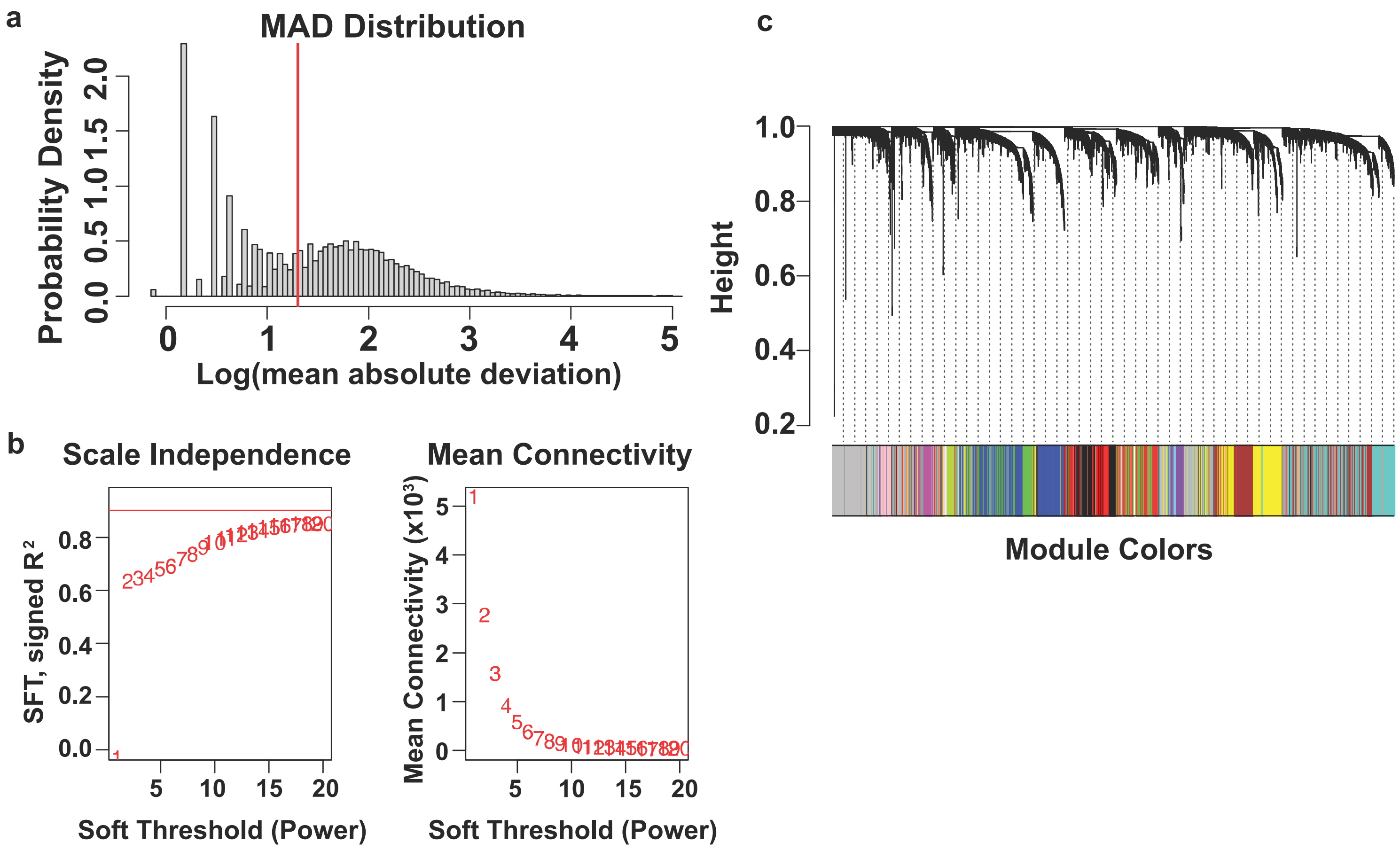

**Supplementary Figure 3. WGCNA of the hepatic transcriptome.** **a**, Mean absolute deviation (MAD) distribution of expressed hepatic transcripts, used to filter low-variance genes prior to network construction. **b**, Scale-free topology fit index (left y-axis) and mean network connectivity (right y-axis) as a function of soft-thresholding power (β), with the selected power indicated by the dashed line. A scale-free topology fit index of $R^2 \geq 0.80$ was used as the threshold for power selection. **c**, Hierarchical clustering dendrogram of all retained hepatic transcripts following soft-thresholding, with module color assignments from the dynamic tree-cut algorithm depicted below. The 19 non-overlapping co-expression modules are each designated by a unique color. Related to Figure 4.

### Supplementary Figure 4

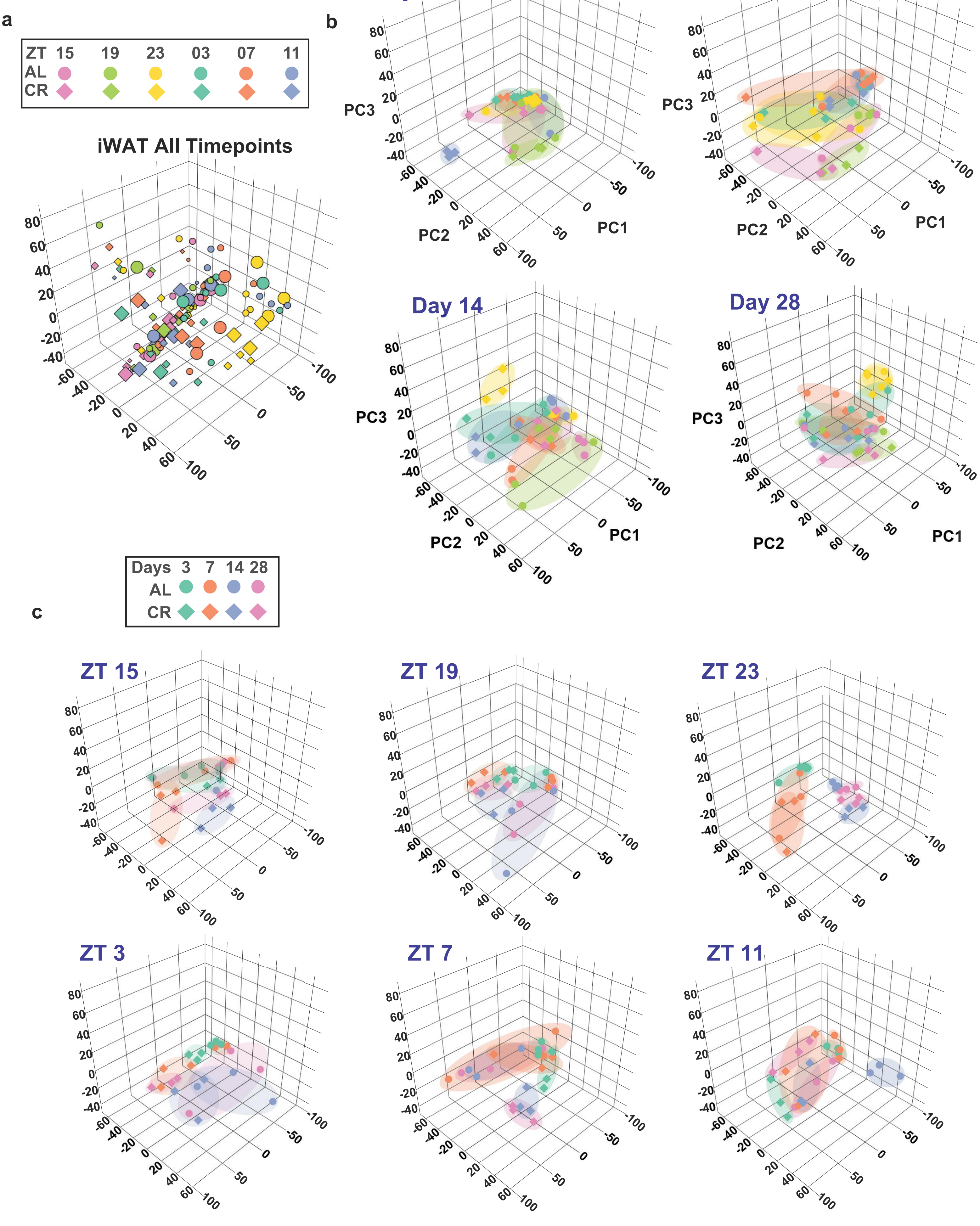

**Supplementary Figure 4. UpSet plots depicting gene intersections across all 19 hepatic co-expression modules. a,** UpSet plots illustrating the overlap of kME-expanded sets (|kME| > 0.6) across all 19 hepatic WGCNA modules at Days 3, 7, 14, and 28. Horizontal bars indicate total gene count per module; vertical bars indicate the size of intersecting gene sets. **b**, Within-module degree z-score and participation coefficient. **Related to Figure 4**.

### Supplementary Figure 5

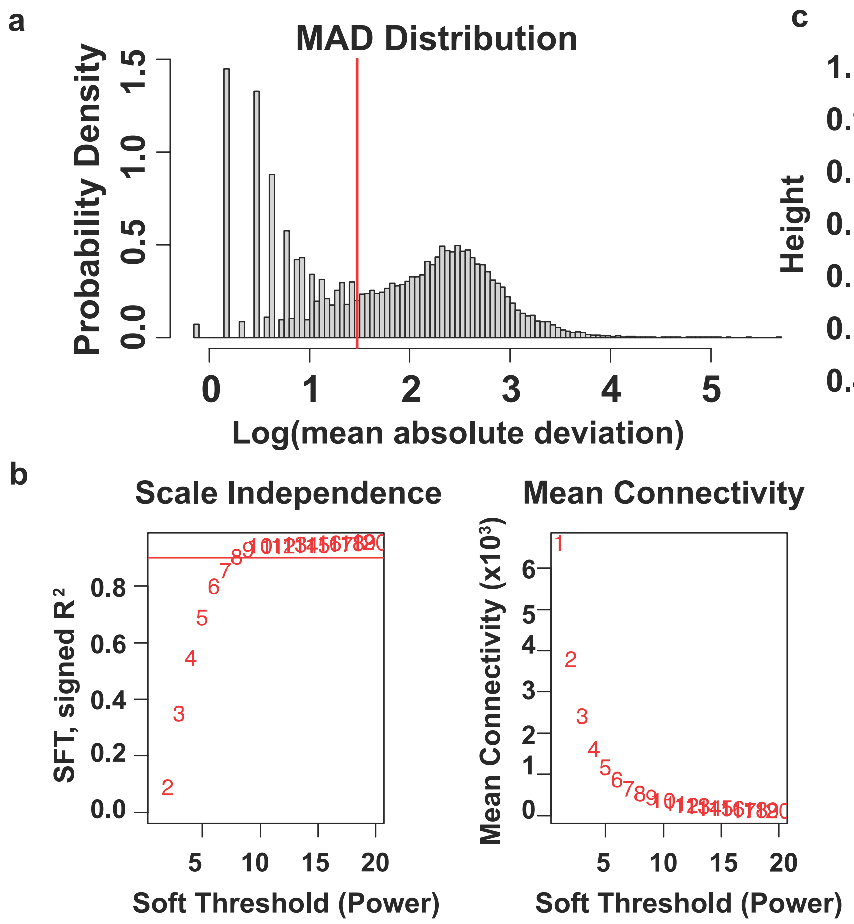

**Supplementary Figure 5. Principal component analysis of the iWAT transcriptome.** **a**, PCA stratified by dietary condition (AL vs. CR). **b**, PCA stratified by adaptive timepoint. **c**, PCA stratified by time of day, with ZT23 exhibiting the strongest separation between AL and CR cohorts. Data represent normalized transcript counts from bulk RNA sequencing. n = X per group. Related to Figure 6.

### Supplementary Figure 6

a

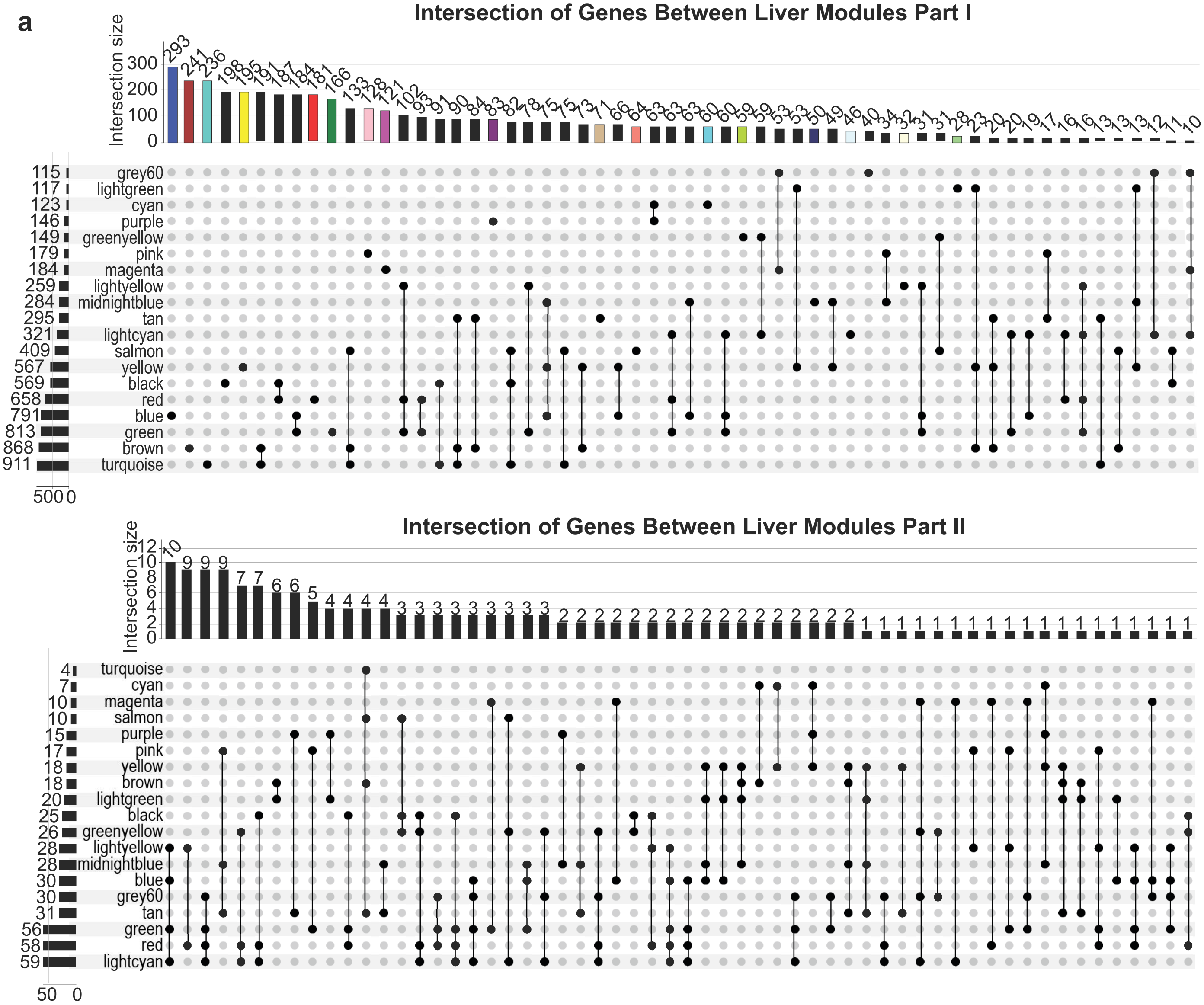


b

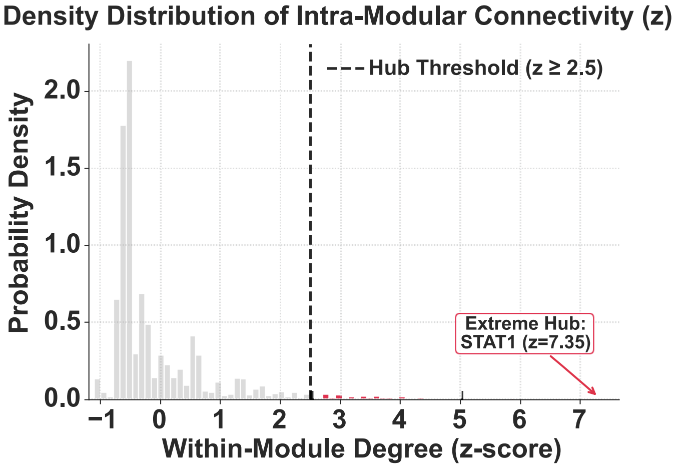

**Supplementary Figure 6. WGCNA of the iWAT transcriptome.** **a**, Mean absolute deviation (MAD) distribution of expressed hepatic transcripts, used to filter low-variance genes prior to network construction. **b**, Scale-free topology fit index (left y-axis) and mean network connectivity (right y-axis) as a function of soft-thresholding power (β), with the selected power indicated by the dashed line. A scale-free topology fit index of $R^2 \geq 0.80$ was used as the threshold for power selection. **c**, Hierarchical clustering dendrogram of all retained hepatic transcripts following soft-thresholding, with module color assignments from the dynamic tree-cut algorithm depicted below. The 11 non-overlapping co-expression modules are each designated by a unique color. **Related to Figure 6**.

### Supplementary Figure 7

a

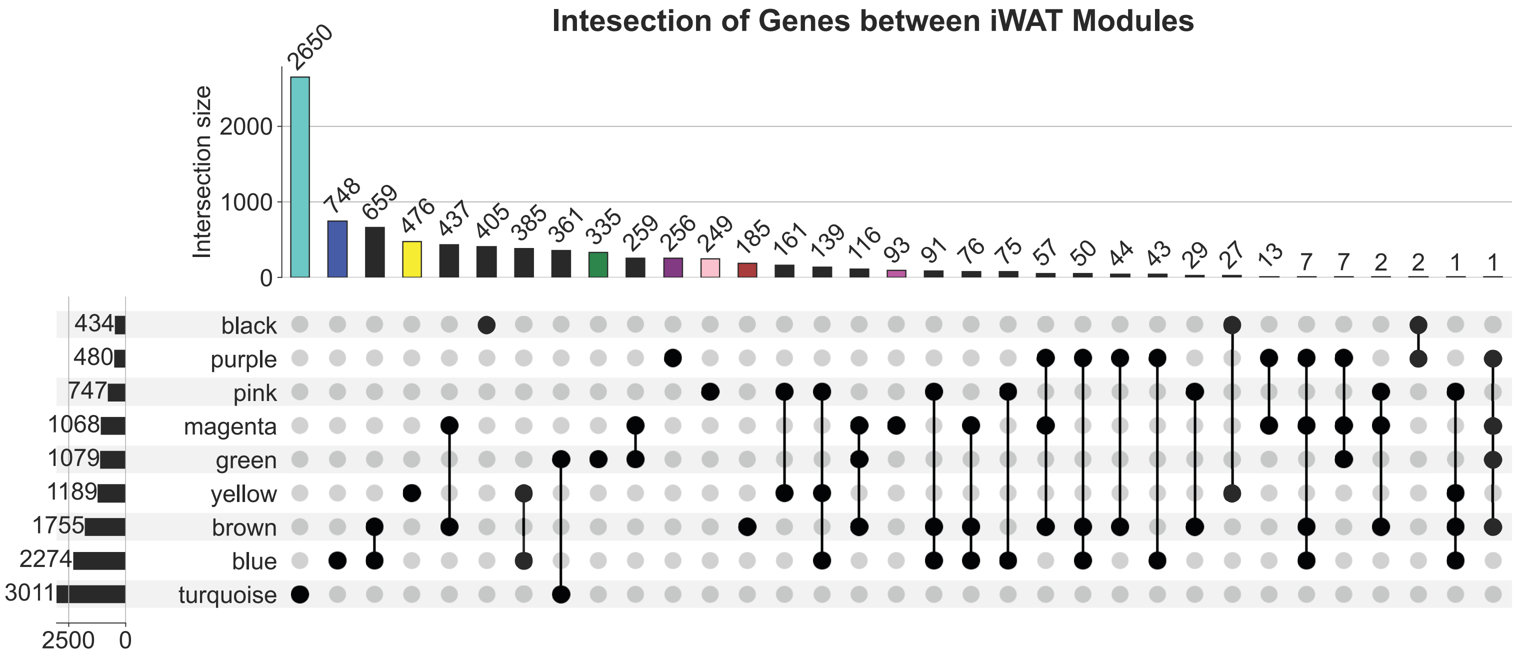


b

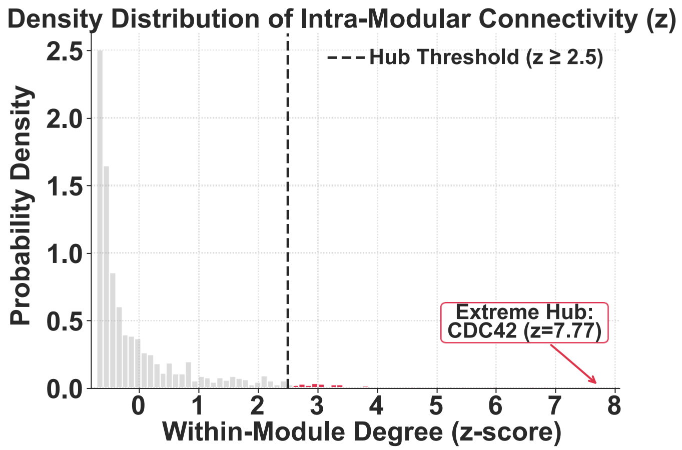

**Supplementary Figure 7. UpSet plots depicting DEG intersections across all 11 iWAT co-expression modules. a,** UpSet plots illustrating the overlap of kME-expanded sets (|kME| > 0.6) across all 11 iWAT WGCNA modules at Days 3, 7, 14, and 28. Horizontal bars indicate total gene count per module; vertical bars indicate the size of intersecting gene sets. **b**, Within-module degree z-score and participation coefficient. **Related to Figure 6**.

#### Supplementary Figure 8

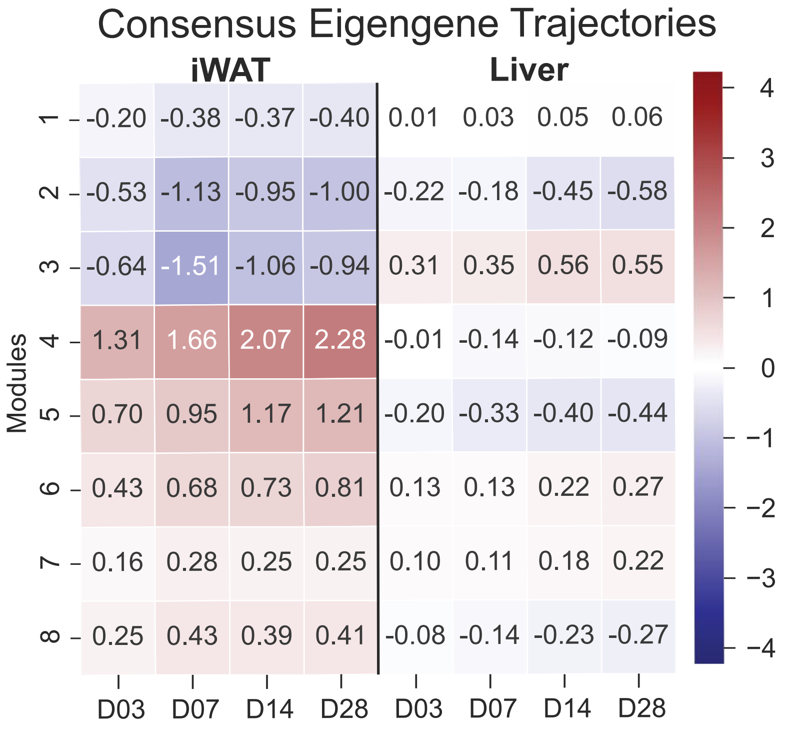

**Supplementary Figure 8.Consensus eigengene profiles for cross-tissue co-expression modules.**

Consensus eigengene values (mean log2 fold-change across all module members) for the eight modules derived from hierarchical clustering of 1,232 transcripts regulated in both liver and iWAT (adjusted $P < 0.05$) across Days 3, 7, 14, and 28, shown separately for each tissue. Summary heatmaps are shown in **Extended Data Figs. 7–8.**
